# Intra- and Inter-Specific Investigations of Skeletal DNA Methylation and Femur Morphology in Primates

**DOI:** 10.1101/554618

**Authors:** Genevieve Housman, Ellen E. Quillen, Anne C. Stone

## Abstract

**Objectives:** Epigenetic mechanisms influence the development and maintenance of complex phenotypes and may also contribute to the evolution of species-specific phenotypes. With respect to skeletal traits, little is known about the gene regulation underlying these hard tissues or how tissue-specific patterns are associated with bone morphology or vary among species. To begin exploring these topics, this study evaluates one epigenetic mechanism, DNA methylation, in skeletal tissues from five nonhuman primate species which display anatomical and locomotor differences representative of their phylogenetic groups.

**Materials and Methods:** First, we test whether intra-specific variation in skeletal DNA methylation is associated with intra-specific variation in femur morphology. Second, we identify inter-specific differences in DNA methylation and assess whether these lineage-specific patterns may have contributed to species-specific morphologies. Specifically, we use the Illumina Infinium MethylationEPIC BeadChip to identify DNA methylation patterns in femur trabecular bone from baboons (n=28), macaques (n=10), vervets (n=10), chimpanzees (n=4), and marmosets (n=6).

**Results:** Significant differentially methylated positions (DMPs) were associated with a subset of morphological variants, but these likely have small biological effects and may be confounded by other variables associated with morphological variation. Conversely, several species-specific DMPs were identified, and these are found in genes enriched for functions associated with complex skeletal traits.

**Discussion:** Overall, these findings reveal that while intra-specific epigenetic variation is not readily associated with skeletal morphology differences, some inter-specific epigenetic differences in skeletal tissues exist and may contribute to evolutionarily distinct phenotypes. This work forms a foundation for future explorations of gene regulation and skeletal trait evolution in primates.

## Introduction

Primates distinguish themselves from other mammals with their unique suite of anatomical features that initially enabled arboreal niche occupation and subsequently evolved to fit a myriad of habitats and forms of locomotion, the most unique being hominin bipedalism. The resulting morphological variation that has evolved across taxa has its foundation in underlying skeletal anatomies. Such skeletal morphologies are used to both characterize extant primate diversity and reconstruct the anatomy and locomotor capabilities of extinct primate species (Ankel-Simons, 2007; Fleagle, 1999). The range of mechanisms that enable the development of skeletal features are not entirely understood, though. Skeletal features related to varied body forms are often described as the result of environmental adaptations. However, skeletal morphology is more accurately defined as the result of complex processes, including environmental, genetic, and epigenetic mechanisms. While environmental (Lewton, 2017; Lewton, Ritzman, Copes, Garland, & Capellini, 2018) and genetic (Joganic et al., 2017; Ritzman et al., 2017) factors have been readily explored, the important roles of epigenetic factors, such as DNA methylation, in skeletal tissue development and maintenance have only recently been identified (Delgado-Calle et al., 2013; Rushton et al., 2014; Simon & Jeffries, 2017).

For instance, epigenetic processes are influential in regulating skeletal muscle development (Palacios & Puri, 2006) which can impact the adjacent skeletal scaffolding system. Several genes involved in human skeletal development appear to be differentially methylated across fetal and adult developmental stages (de Andrés et al., 2013). Lastly, methylation variation in humans and model organisms has been implicated in several skeletal pathologies and disorders, such as osteoporosis and osteoarthritis (Delgado-Calle et al., 2013; den Hollander et al., 2014; Jeffries et al., 2016; Moazedi-Fuerst et al., 2014; Morris et al., 2017; Ostanek, Kranjc, Lovšin, Zupan, & Marc, 2018; Rushton et al., 2014; Simon & Jeffries, 2017). Some of these studies are the first to assess methylation patterns in human skeletal tissues, and such steps are crucial for identifying the relationship between epigenetic variation and skeletal phenotypic variation.

The contributions of epigenetics to primate phenotypic variation were first considered by King and Wilson (1975), who proposed that anatomical and behavioral differences between humans and chimpanzees were more likely “based on changes in the mechanisms controlling the expression of genes than on sequence changes in proteins” (King & Wilson, 1975). Studies to understand methylation variation across species began soon afterwards (Gama-Sosa et al., 1983). General changes to mammalian epigenomes have been examined (Sharif, Endo, Toyoda, & Koseki, 2010), but most epigenetics work in primates has focused on humans – how it varies across distinct tissues within individuals (Slieker et al., 2013), across different individuals (Petronis et al., 2003), across populations (Heyn et al., 2013), in relation to aging processes (Fraga et al., 2005), and in relation to diet (Shelnutt et al., 2004), as well as how it is inherited across generations (Flanagan et al., 2006). These studies have identified interesting intra-specific methylation variation present in humans.

Similarly, epigenetic variation has been identified among primate species. Inter-specific variation in epigenetic signatures was initially inferred from underlying genomic sequences (Bell et al., 2012). For instance, several promoter CpG densities vary across primates. These likely relate to regulatory methylation differences across species as primate CpG densities correlate with methylation levels (Weber et al., 2007). Additionally, gene expression studies, which primarily focus on brain tissues (Babbitt et al., 2010; Cáceres et al., 2003) and a small set of other soft tissues (Blake et al., 2017; Blekhman, Oshlack, Chabot, Smyth, & Gilad, 2008; Karere et al., 2013; Pavlovic, Blake, Roux, Chavarria, & Gilad, 2018; Tung, Zhou, Alberts, Stephens, & Gilad, 2015), have also noted regulatory differences across species. Methylation differences in brain tissues have evolved across primates and contributed to resultant brain phenotypes and disease vulnerabilities (Enard et al., 2004; Farcas et al., 2009; Madrid, Chopra, & Alisch, 2018; Mendizabal et al., 2016; Provencal et al., 2012; Zeng et al., 2012). Thus, methylation-phenotype relationships can be identified in primates. Primate methylation patterns in blood cells and other soft tissues have also been studied, but not to the same degree (Fukuda et al., 2013; Gao et al., 2017; Hernando-Herraez et al., 2013; Lea, Altmann, Alberts, & Tung, 2016; Lindskog et al., 2014; Martin et al., 2011; Molaro et al., 2011; Pai, Bell, Marioni, Pritchard, & Gilad, 2011; Vilgalys, Rogers, Jolly, Mukherjee, & Tung, 2018).

Interestingly, two studies using soft tissues and blood identified differential methylation and expression of genes essential for skeletal development (*RUNX1, RUNX3*, and *COL2A1*) between some primates (Hernando-Herraez et al., 2013; Lindskog et al., 2014). Additionally, the emerging field of ancient epigenetics, which reconstructs methylation patterns from ancient DNA degradation patterns in hominin remains (Smith, Monroe, & Bolnick, 2015; Gokhman et al., 2014; Gokhman, Meshorer, & Carmel, 2016; Gokhman et al., 2017), have found that other skeletal developmental genes (*HOXD* complex) are differentially methylated among modern humans and ancient hominins (Gokhman et al., 2014). These findings suggest that primates do exhibit distinct epigenetic patterns and that primate skeletal epigenetics is an important area of research for uncovering developmental and evolutionary patterns relevant for complex skeletal traits, such as anatomy, locomotion, and disease. Despite these potential avenues of insight, studies of nonhuman primate (NHP) skeletal epigenetics are limited (Housman, Havill, Quillen, Comuzzie, & Stone, 2018).

Overall, there are clear knowledge gaps in our understanding of NHP skeletal complexity in relation to epigenetic variation and epigenetic differences between phylogenetically diverse NHP species. The present study begins to remedy this by assessing how genome-wide and gene-specific DNA methylation in primate skeletal tissues varies intra- and inter-specifically and in relation to femur form. Specifically, for this study, we explored the evolution of the epigenome and its relation to nonpathological skeletal traits by identifying DNA methylation patterns in femur trabecular bone from baboons, macaques, vervets, chimpanzees, and marmosets and assessing intra- and inter-specific methylation variation and its relation to morphology. With this study design, we aimed to assess whether skeletal methylation variation within species is associated with skeletal morphology variation, to what degree skeletal methylation varies among species, and whether species-specific changes in methylation further inform our understanding of skeletal morphology evolution. Additionally, this dataset expands the number of characterized skeletal methylation patterns in NHPs for future primate skeletal epigenetics investigations.

## Materials and Methods

### Ethics Statement

NHP tissue samples included were opportunistically collected at routine necropsy of these animals. No animals were sacrificed for this study, and no living animals were used in this study. Chimpanzee tissues were collected opportunistically during routine necropsy prior to the September 2015 implementation of Fish and Wildlife Service rule 80 FR 34499.

### NHP Samples

NHP samples come from captive colonies of chimpanzees (*Pan troglodytes*), baboons (*Papio spp*.), rhesus macaques (*Macaca mulatta*), and marmosets (*Callithrix jacchus*) from the Southwest National Primate Research Center in Texas, as well as vervets (*Chlorocebus aethiops*) from the Wake Forest/UCLA Vervet Research Colony in North Carolina. Femora were opportunistically collected at routine necropsy of these animals and stored in -20°C freezers at the Texas Biomedical Research Institute after dissection. These preparation and storage conditions ensured the preservation of skeletal DNA methylation patterns. Samples include baboons (n=28), macaques (n=10), vervets (n=10), chimpanzees (n=4), and marmosets (n=6). Age ranges span adulthood for each species and are comparable between each group (Figure 1, Table S1). Scaled ages were also calculated for each sample (Table S1) using published life expectancies for each species (Rowe, 1996). Both sexes are represented (female: n=33, male: n=24, unknown: n=1).

**Figure 1.**
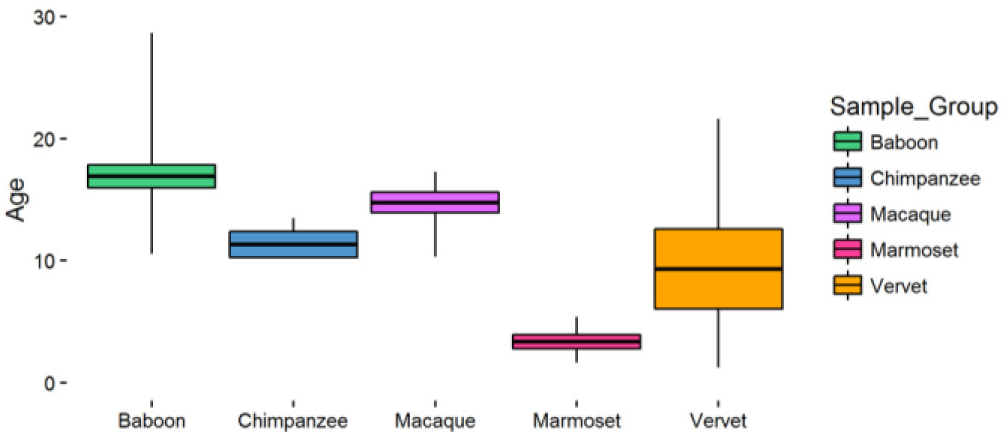
Nonhuman Primate Sample Set Ages. Boxes represent one standard deviation from the average age, and whiskers depict the full range of ages for each species. Baboons (n=28) are 16.90±5.02 years, chimpanzees (n=4) are 11.31±1.87 years, macaques (n=10) are 14.75±2.65 years, marmosets (n=6) are 3.34±1.41 years, and vervets (n=10) are 9.31±10.30 years. Additional details can be found in Table S1.

### Assessment of Femur Morphologies

On the right femora of NHP samples, 29 linear morphology traits (Figure 2, Table S6) were measured using calipers. These measurements characterize overall femur shape (McHenry & Corruccini, 1978; Terzidis et al., 2012). All measurements were collected by one researcher, and intra-observer error for each measurement was determined by performing triplicate measurements on approximately 10% of the samples in each species. These measurements were spaced throughout the entire data collection period. Error was calculated as the mean absolute difference divided by the mean (Corner, Lele, & Richtsmeier, 1992; White & Folkens, 2000). All measurements that were retained for downstream analyses had errors of less than 5%, and the only measurement excluded was macaque intercondylar notch depth (error = 6.62%) (File S2).

**Figure 2.**
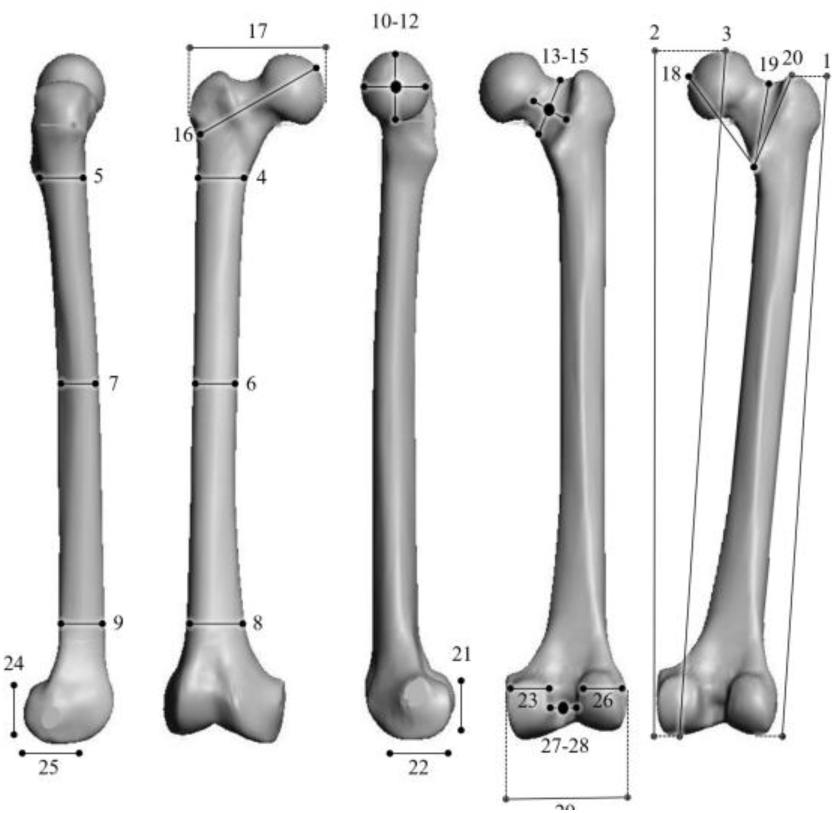
Nonhuman Primate Morphological Measurements. See Table S6 for a detailed description of these measurements.

### Genome-Wide DNA Methylation Profiling

Trabecular bone cores were obtained from the medial condyles on the right distal femora of each NHP sample using a drill press that cored transversely through the condyle leaving the articular surface preserved (Supplemental Text). Cortical bone was removed from cores using a Dremel, and the remaining trabecular bone cores were pulverized into bone dust using a SPEX SamplePrep Freezer/Mill. DNA was extracted from femoral trabecular bone using a phenol-chloroform protocol optimized for skeletal tissues (Barnett and Larson 2012). Genome-wide DNA methylation was assessed using Illumina Infinium MethylationEPIC microarrays (EPIC array) (Supplemental Text). The array data discussed in this publication have been deposited in NCBI’s Gene Expression Omnibus and are accessible through GEO SuperSeries accession number GSE103332, which includes the SubSeries accession numbers GSE103279, GSE103271, GSE103280, GSE94677, GSE103328, and GSE103287.

### Methylation Data Processing

Using previously described methods (Housman et al., 2018), raw EPIC array data were normalized and converted to β values which represent the average methylation levels at each site (0 = completely unmethylated sites, 1 = fully methylated sites), and M values which are the log transformed ratio of methylated signal to unmethylated signal. Probes with failed detection levels (p-value > 0.05) in greater than 10% of samples and samples with greater than 30% of probes with detection level failures were removed from downstream analyses. Using previously described methods (Housman et al., 2018), probes with sequence mismatches to NHP genomes, which could produce biased methylation measurements, were computationally filtered out and excluded from downstream analyses (Supplemental Text, Figures S1-S3, Tables S2-S5, File S1). Additionally, cross-reactive probes (McCartney et al., 2016), probes containing SNPs at the CpG site, probes detecting SNP information, probes detecting methylation at non-CpG sites, and probes targeting sites within the sex chromosomes were removed (Aryee et al., 2014; Fortin, Triche, & Hansen, 2016). Lastly, we assessed whether NHP SNPs overlapped with targeted probes (Supplemental Text, Table S2, File S1). The resulting finalized datasets (Figure S4) were then used in further statistical analyses.

### Statistical Analysis of Differential Methylation

In order to identify sites that were significantly differentially methylated across comparative groups, we designed and tested general linear models (GLMs) which related the variables of interest (morphological measures and species membership) to the DNA methylation patterns for each site, while accounting for the effects of additional variables, batch effects, and latent variables (Maksimovic, Phipson, & Oshlack, 2016). Sites found to have significant associations were classified as significant differentially methylated positions (DMPs). Overall, GLMs were used to estimate differences in methylation levels associated with femur morphologies within each taxonomic group (intra-specific) and between each taxonomic group (inter-specific). Specific details about each set of GLMs are provided in the following sections with additional details in the Supplemental Text.

### Intra-Specific Analyses

For the intra-specific analyses, variables included in each GLM were the femur morphologies within each taxonomic group, sex, age, and steady state weight when known, as well as unknown latent variables calculated using the iteratively re-weighted least squares approach in the sva package in R (Jaffe & Irizarry, 2014; Leek, Johnson, Parker, Jaffe, & Storey, 2012; Leek & Storey, 2007, 2008). Latent variables estimated for each morphology were included to help mitigate any unknown batch and cell heterogeneity effects on methylation variation at each site (Table S7). Each GLM design matrix was fit to corresponding M value array data by generalized least squares using the limma package in R (Huber et al., 2015; Phipson, Lee, Majewski, Alexander, & Smyth, 2016; Ritchie et al., 2015), and the estimated coefficients and standard errors for each morphology were computed. Lastly, for each coefficient, an empirical Bayes approach was applied using the limma package in R (Lönnstedt & Speed, 2002; McCarthy & Smyth, 2009; Phipson, Lee, et al., 2016; Smyth, 2004) to compute moderated t-statistics, log-odds ratios of differential methylation, and associated p-values adjusted for multiple testing (Benjamini & Hochberg, 1995). Significant DMPs for the effect of each morphology were defined as those having log fold changes in M values corresponding to an adjusted p-value of less than 0.05. Lastly, the gene ontology (GO) and KEGG pathway enrichment for significant CpGs was determined using the missMethyl package in R (Benjamini & Hochberg, 1995; Geeleher et al., 2013; Phipson, Maksimovic, & Oshlack, 2016; Ritchie et al., 2015; Young, Wakefield, Smyth, & Oshlack, 2010), which takes into account the differing number of probes per gene present on the array.

### Inter-Specific Analyses

For the inter-specific analyses, variables included in the GLM were taxonomic grouping, sex, age, known batch effects (e.g., array number and position), and unknown latent variables calculated using the method described above. An additional GLM which replaced the age variable with ages scaled to species-specific life expectancies (Rowe, 1996) was also tested. The six latent variables estimated were included to help mitigate any unknown batch and cell heterogeneity effects on methylation variation at each site. The GLM design matrix was fit to the M value array data using the method described above, and the estimated coefficients and standard errors for taxonomic group effects were computed. As described above, moderated t-statistics, log-odds ratios of differential methylation, and associated p-values adjusted for multiple testing were computed, and significant DMPs for the effect of taxonomy were defined as those having log fold changes in M values corresponding to an adjusted p-value of less than 0.05.

To determine only those methylation differences that represent fixed changes between genera, we used methods similar to those described in (Hernando-Herraez et al., 2013). Briefly, significant DMPs were identified between all possible pairwise comparisons of taxa (n=10: baboon-macaque, baboon-vervet, baboon-chimpanzee, baboon-marmoset, macaque-vervet, macaque-chimpanzee, macaque-marmoset, vervet-chimpanzee, vervet-marmoset, chimpanzee-marmoset). A significant DMP was then defined as taxon-specific if it was found to be significant in all four pairwise comparisons containing the taxon of interest but not found in any of the remaining pairwise comparisons. The GO and KEGG pathway enrichment of these DMPs was then determined as described above.

Additionally, global changes in methylation were calculated using distance matrices (Hernando-Herraez et al., 2013) of the methylation levels for all finalized 39,802 filtered probes. These changes were assessed at a species-level by averaging the β values per probe within each species. We then used Euclidean distances to calculate the difference between every two species. Neighbor joining trees were estimated from these distances using the ape package in R (Paradis, Claude, & Strimmer, 2004). For each resulting tree, 1000 bootstraps were performed to determine confidence values for each branch. Global changes in methylation were also assessed at the individual-level using Euclidean distances to calculate the difference between every two individuals.

### DNA Methylation Profiling and Analyses of HOXD10

Based on the inter-specific DNA methylation patterns identified in this study and those identified in other evolutionary anthropological studies (Gokhman et al. 2014), the *HOXD10* gene was selected for subsequent DNA methylation profiling and analysis at a higher resolution using gene-specific sequencing techniques. Specifically, primers were designed and optimized to PCR amplify regions spanning across the entire *HOXD10* gene, as well as upstream and downstream several hundred bases (hg19 chr2:176980532-176985117), in each NHP species for regular and bisulfite treated DNA (Tables S14-S17). All gene-specific assays were performed in a subset of the samples tested using the EPIC array and included chimpanzees (n=3), baboons (n=3), macaques (n=3), vervets (n=3), and marmosets (n=3) (Table S1). Additionally, a subset of these assays that targeted the region around one test locus (cg02193236) were performed in all of the samples (Table S1). As described above, DNA was extracted from femoral trabecular bone using a phenol-chloroform protocol optimized for skeletal tissues (Barnett and Larson 2012). DNA was bisulfite converted using the EZ DNA MethylationTM Gold Kit according to the manufacturer’s instructions (Zymo Research). Successful PCR amplification was confirmed using gel electrophoresis. Gene-specific PCR products were then purified using an exonuclease I and shrimp alkaline phosphatase protocol and sequenced on the Applied Biosystems 3730 capillary sequencer at the DNA Laboratory at Arizona State University.

Regular and bisulfite sequences were aligned to the appropriate NHP references within the Enredo-Pecan-Orthus (EPO) whole-genome multiple alignments of several primate genomes [Ensembl Compara.8_primates_EPO] (Paten, Herrero, Beal, et al. 2008; Paten, Herrero, Fitzgerald, et al. 2008) using MEGA7 (Kumar et al. 2016). Manual annotation of these sequences within each sample confirmed that the gene sequences belong to the appropriate primate species and that the regular and bisulfite treated sequences only differ in cytosine composition. The number and distribution of methylated loci throughout the *HOXD10* gene were then identified and compared within and among species to provide a higher resolution of methylation variation within this targeted gene. Lastly, with respect to the test locus (cg02193236) evaluated in all samples, methylation levels across this region were compared with corresponding methylation levels of this region as determined using the EPIC array.

## Results

The aim of this study was to identify DNA methylation patterns in skeletal tissues from several NHP species in order to determine how skeletal methylation varies at different taxonomic scales and in relation to complex skeletal traits. We evaluated DNA methylation patterns in femur trabecular bone from five NHP species – baboons, macaques, vervets, chimpanzees, and marmosets (Figure 1, Table S1). We used the Illumina Infinium MethylationEPIC BeadChip (EPIC array), which was determined to be effective at identifying DNA methylation patterns in NHP DNA (Supplemental Text, Figures S1-S4, Tables S2-S5, File S1). Further, we selected probes appropriate for intra- and inter-specific comparison, which removed the effect of sequence differences among individuals and species as a reason for methylation differences. With these data, we first tested whether intra-specific variation in skeletal DNA methylation is associated with intra-specific variation in femur morphology. Second, we identified inter-specific differences in DNA methylation and assessed whether these lineage-specific patterns may have contributed to species-specific morphologies.

### Genome-Wide Intra-Specific Differential Methylation and Morphological Variation

Measurements of 29 linear morphology traits (Figure 2, Table S6) were collected from each NHP right femur. All measurements had less than 5% error, except those for intercondylar notch depth in macaques (Figure S5, File S2). Significant DMPs associated with each intra-specific linear morphology were interrogated from 189,858 sites in in baboons, 190,898 sites in macaques, 191,639 sites in vervets, 576,804 sites in chimpanzees, and 68,709 sites in marmosets (Tables S7-S8). In baboons (n=28), 1 DMP was hypomethylated with increasing bicondylar femur length and increasing maximum femur length. In macaques (n=10), 1 DMP was hypermethylated with increasing proximal femur width, 1 DMP was hypermethylated with increasing medial condyle width, and 6 DMPs were hypomethylated with increasing medial condyle width. In vervets (n=10), 1 DMP was hypomethylated with increasing superior shaft width, 2 DMPs were hypomethylated with increasing inferior shaft width, and 1 DMP was hypermethylated with increasing anatomical neck height. In chimpanzees (n=4), 216 DMPs were hypomethylated and 57 DMPs were hypermethylated with increasing anatomical neck length. Lastly, in marmosets (n=6), no DMPs were associated with morphological variation (Table S9, File S3).

While the maximum absolute change in mean methylation (Δβ) for most of these DMPs is greater than 10% (Δβ = 0.1), the change in methylation between the individual with the largest morphology measurement and the individual with the smallest morphology measurement is less than 0.1 for several DMPs (File S3). Tests for enrichment of gene ontology (GO) and KEGG pathway functions were done for the 3 intra-specific morphologies that had more than 2 DMPs associated with them. However, no GO biological processes were found to be enriched in DMPs associated with either macaque medial condyle width or chimpanzee anatomical neck length. Additionally, KEGG pathway functions were only found to be enriched in DMPs associated with chimpanzee anatomical neck length, and these pathways were predominantly involved in immune system cell signaling and differentiation (Table S10).

### Genome-Wide Inter-Specific Differential Methylation

To determine how methylation varies inter-specifically, differential methylation was interrogated from the 39,802 probes that were shared among NHP species (Table S11, File S4). Species-specific DMPs were determined by identifying DMPs that were significant in all 4 pairwise comparisons containing the taxon of interest but not in any of the remaining pairwise comparisons. These methods identified 650 species-specific DMPs in baboons, 257 in macaques, 639 in vervets, 2,796 in chimpanzees, and 13,778 in marmosets (Table S11). Similar results were obtained using a model that accounted for scaled age based on life expectancy instead of exact age (660 baboon-specific DMPs, 260 macaque-specific DMPs, 651 vervet-specific DMPs, 2,720 chimpanzee-specific DMPs, and 13,689 marmoset-specific DMPs) (File S4). Thus, only the species-specific DMPs identified using the model that accounted for exact age were used in downstream analyses. Additionally, given the disproportionate number of marmoset-specific DMPs identified, caution should be used when interpreting the biological relevance of these data. Comparably, methylation patterns also distinguish broader taxonomic groups, such as cercopithecoids (baboons, macaques, and vervets), apes (chimpanzees), and platyrrhines (marmosets). Specifically, 2,655 DMPs were found to be specific to cercopithecoids, 1,869 were found to be ape-specific, and 14,985 were found to be specific to platyrrhines (File S5). It should be noted that the ape and platyrrhine categories only include one taxon each – chimpanzee and marmoset, respectively. Similar results were obtained using a model that accounted for scaled age based on life expectancy instead of exact age (2,660 cercopithecoid-specific DMPs, 1,797 ape-specific DMPs, and 14,963 platyrrhine-specific DMPs) (File S5).

Species-specific DMPs spanned 7,320 genes with an average of 2.2±2.0 (s.d.) significant probes per gene (Table S12). Over half of these genes contain just 1 significant species-specific DMP (n=3,816), and a portion of these are due to only 1 EPIC array probe targeting each gene (n=1,788). The remaining genes that contain at least 2 significant species-specific DMPs (n=3,503) have an average of 3.4±2.3 (s.d.) significant probes per gene (range: 2-34). Additionally, these species-specific DMPs covered a range of locations with respect to genes and CpG islands (Table S12), indicating that these species-specific changes in methylation are distributed throughout the genome. Using various Δβ cutoff thresholds decreases the final number of species-specific DMPs to varying degrees (Table S11, File S4). Counter to these decreases in numbers, though, species-specific DMPs cover a range of locations with respect to genes and CpG islands regardless of the Δβ cutoff threshold (Table S12).

Overall, across Δβ cutoff thresholds, more species-specific DMPs were found in the platyrrhine marmosets, followed by the great ape chimpanzees, and lastly the cercopithecoid baboons, macaques, and vervets. Additionally, the proportions of hypermethylated and hypomethylated species-specific DMPs within each taxon remain fairly constant across Δβ cutoff thresholds (Table S11). In baboons, macaques, vervets, and chimpanzees, more than half of all species-specific DMPs show patterns of hypermethylation, and in marmosets, more than half of all species-specific DMPs show patterns of hypomethylation. The only disruption in these trends is in chimpanzees when no Δβ threshold is applied. Nevertheless, species-specific DMPs with different Δβ cutoff thresholds do differ in their abilities to cluster animals into taxonomic groups (Figure S6, File S4). For all thresholds, apes, cercopithecoids, and platyrrhines cluster into distinct groups. However, within cercopithecoids, vervets only cluster into a distinct species group with a Δβ ≥ 0.2 threshold, and the baboon-macaque clade require a Δβ ≥ 0.3 threshold. Lastly, species-specific clustering of baboons, macaques, and vervets only occurs with a Δβ ≥ 0.4 threshold (Figure 3).

**Figure 3.**
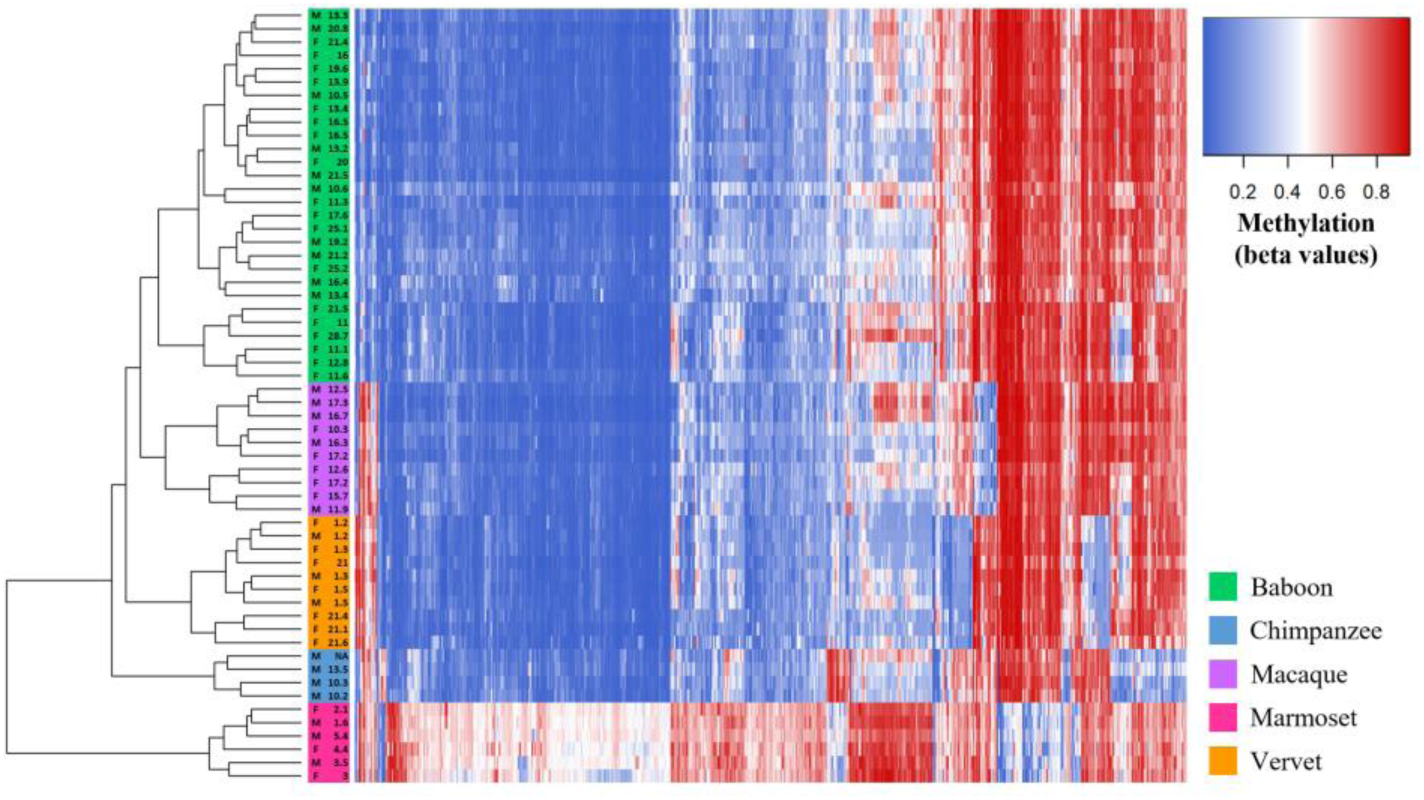
Methylation Levels at Species-Specific DMPs with Δβ≥0. 4 Identified in the Inter-Specific Study. Heatmap depicting the DNA methylation levels (β values) of all species-specific DMPs with average absolute Δβ values greater than 0.4 between each taxonomic group (x-axis) in all nonhuman primate samples (n=58). The sex and age of each nonhuman primate are also provided (y-axis). Red indicates higher methylation at a DMP, while blue indicates lower methylation at a DMP. The dendrogram of all samples (y-axis) clusters individuals based on the similarity of their methylation patterns. Samples cluster based on species-level taxonomic groupings and as predicted based on known species phylogenetic histories.

Additionally, global changes in methylation across all 39,802 probes were evaluated. Average global changes within each species reveal that apes, cercopithecoids, and platyrrhines are phylogenetically distinct from one another, and these divergences are well supported (Figure 4). Similarly, when phylogenetic relationships are evaluated using the global changes in methylation of individual animals, distinct lineages are again formed between apes, cercopithecoids, and platyrrhines (Figure S7). However, within the cercopithecoid clade, several poorly supported branches result in baboons, macaques, and vervets not forming distinct lineages. Phylogenetic separation of these cercopithecoid species into distinct lineages is only possible when the methylation changes considered are reduced to only include species-specific DMPs with a Δβ ≥ 0.4 threshold (Figure S8).

**Figure 4.**
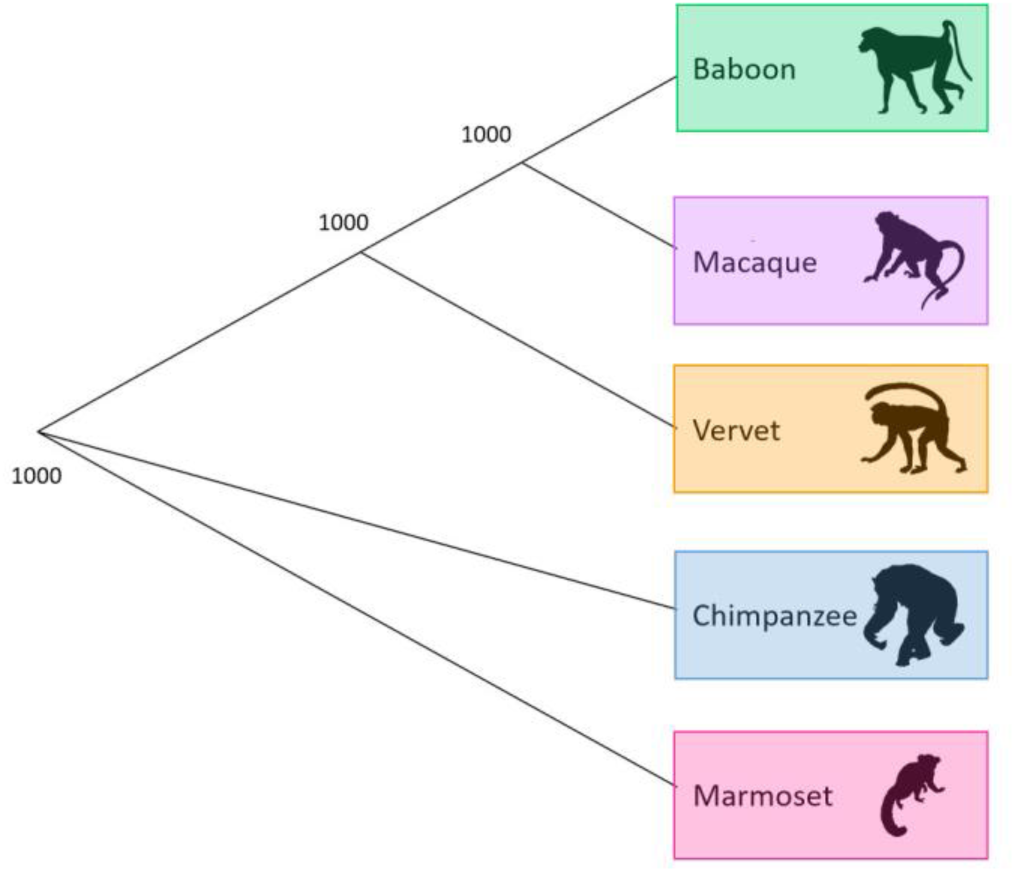
Phylogeny Based on Average Species-Level Global Changes in Methylation. Observed phylogenetic relationship among nonhuman primates when considering average species-level global changes in methylation. This tree was constructed using the methylation levels for all finalized 39,802 filtered probes. We averaged the β values per probe within a species, used Euclidean distances to calculate the difference between every two species, and estimated a neighbor joining tree using this distance matrix. For the resulting tree, 1000 bootstraps were performed to determine confidence values for each branch. The number provide at each node indicates the number of bootstrap replicates that support it out of the 1000 performed.

Species-specific DMPs also show associations with genes that have a wide array of GO biological processes (File S6) and KEGG pathway functions (File S7). Cellular adhesion is a primary GO function found to be highly enriched in species-specific DMPs from baboons, macaques, vervets, chimpanzees, and marmosets. Species-specific DMPs for chimpanzees and marmosets are also enriched for genes involved in the regulation of transcription and gene expression. Additionally, enrichment of genes involved in anatomical developmental processes is found in baboons, vervets, chimpanzees, and marmosets. Chimpanzees and marmosets show further enrichment of genes contributing to pattern specification processes, limb development, and skeletal system development. Moreover, marmoset species-specific DMPs are enriched for genes with functions very closely related to skeletal development, such as osteoblast differentiation and ossification, as well as genes involved in metabolism and the development of other organ systems including skeletal muscles, nerves, the brain, the heart, blood vessels, kidneys, eyes, and ears (File S6). Several enriched pathways reinforce these molecular functions, and additional pathways related to cancers and other disease were also identified (File S7).

Lastly, out of the species-specific DMPs identified, some were found to overlap with those previously identified as being differentially methylated among primates. These included genes within the *HOXD* cluster which have been found to be differentially methylated among modern and ancient hominins (Gokhman et al., 2014). Specifically, 1 baboon-specific DMP is hypermethylated in *HOXD9*, and 2 chimpanzee-specific DMPs are hypomethylated in *HOXD9* (File S4). Additional genes in the *HOXD* cluster (*HOXD8* and *HOXD10*) contain marmoset-specific DMPs, and genes such as *ARTN, COL2A1*, and *GABBR1* which have been found to be differentially methylated among modern humans and great apes (Hernando-Herraez et al., 2013), also contain marmoset-specific DMPs (File S4). However, given the unusual nature of the marmoset data, these results will not be discussed further.

### Inter-Specific DNA Methylation Profiling of HOXD10

Based on the inter-specific DNA methylation patterns identified in this study (Figure 5, Table S13) and those identified in other evolutionary anthropological studies (Gokhman et al. 2014), the *HOXD10* gene was selected for subsequent DNA methylation profiling and analysis at a higher resolution using gene-specific sequencing techniques. Regular and bisulfite sequences of several regions in the *HOXD10* gene were generated (Tables S14-S17). First, loci across the entire *HOXD10* gene were examined from a subset of EPIC array samples – 3 baboons, 3 macaques, 3 vervets, 3 chimpanzees, and 3 marmosets (Figure 6, Tables S1 and S18, File S8-S9). Additionally, one test locus (cg02193236) and its surrounding region in *HOXD10* were surveyed in all NHP samples in order to confirm the reliability of the EPIC array in assessing DNA methylation levels (Figure S9, Tables S1 and S19, File S10-S11). This test, which showed similar methylation patterns to those identified with the EPIC array, supplemented previous methylation array replication studies that have found resulting data to be highly correlated in replicate chimpanzee tissue samples (>98% correlated) (Pai et al., 2011) and in replicate macaque tissue samples (97% correlated) (Ong et al., 2014). Following the alignment of these sequences to the appropriate NHP references, the presence and absence of methylation across the *HOXD10* gene in each animal was determined (Tables S18-S19, Files S8-S11). These data reveal that across the *HOXD10* gene, NHPs display generally low methylation with some clustered increased amounts of methylation upstream of the gene and at the start of the gene body (Figure 6). Additionally, in marmosets, the bisulfite sequence findings are consistent with methylation patterns identified using the EPIC array, so these data are unable to further clarify the unusual distribution of methylation levels discovered in these samples (Figure S4).

**Figure 5.**
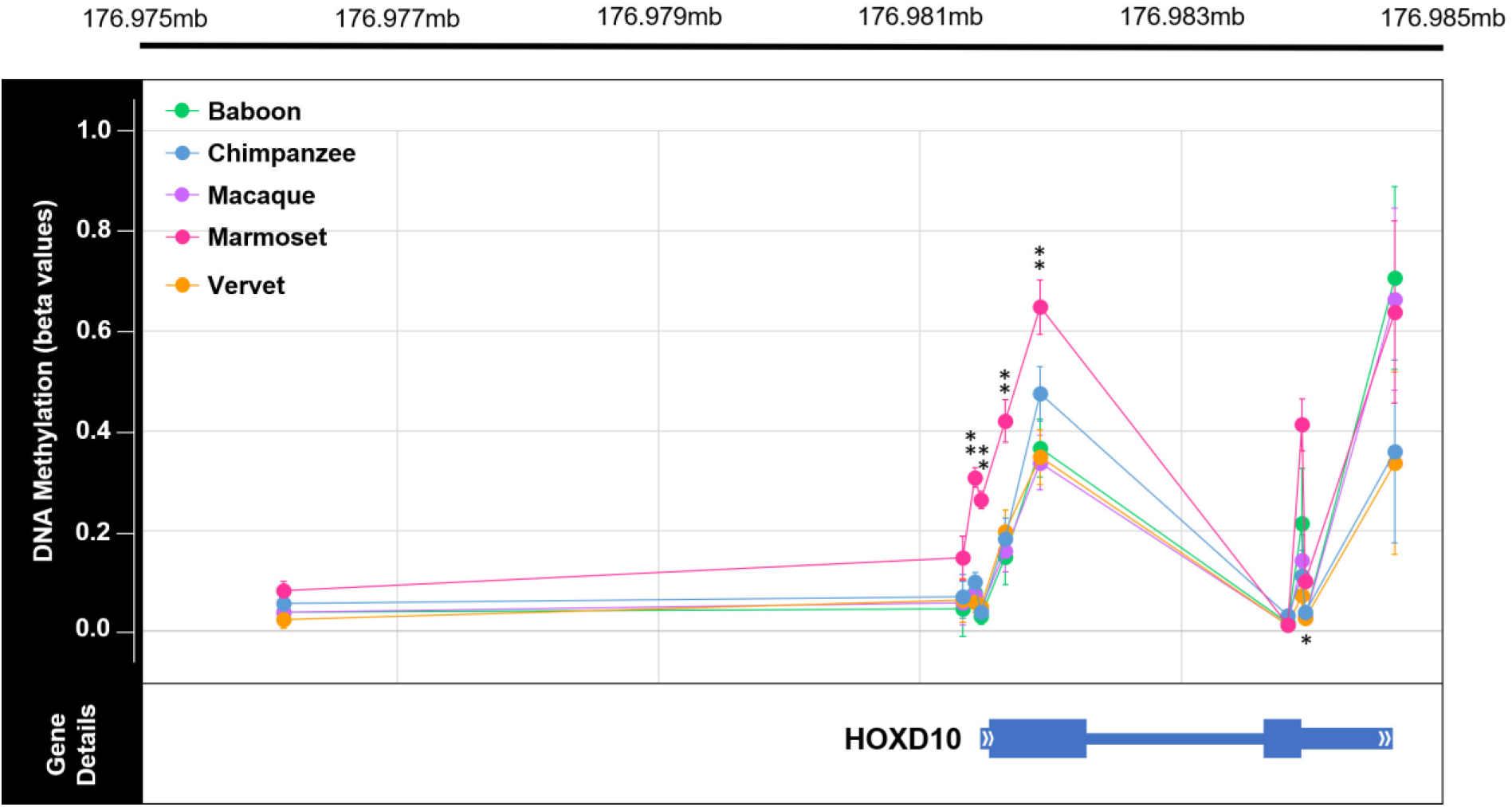
Genome-Wide Methylation Levels Across *HOXD10* in Nonhuman Primates. Plot of the methylation levels of significant DMPs across the *HOXD10* gene (hg19 chr2:176981492-176984670). Plot shows the average β values for each DMP with error bars indicating 1 standard deviation in each direction for each comparative group (teal = baboon, orange = chimpanzee, purple = macaque, pink = marmoset, and light green = vervet). DMP chromosomal position in relation to the *HOXD10* gene is also depicted. This gene is of interest because it has been found to be differentially methylated in ancient and modern hominin species (Gokhman et al., 2014). Of the sites depicted here, 5 DMPs were found to show significant species-specific methylation in marmosets. Of the 5 species-specific DMPs in the *HOXD10* gene of marmosets, 4 have Δβ between 0.2 and 0.3 (**) and 1 has a Δβ < 0.1 (*). See Table S13 for additional information. In the *HOXD10* gene, the two exons are denoted with the thickest bars (exon 1: hg19 chr2:176981561-176982306; exon 2: hg19 chr2:176983681-176983959), the UTRs are denoted with bars of intermediate thickness (5’UTR: hg19 chr2:176981491-176981561; 3’UTR: hg19 chr2:176983959-176984670), and the one intron is denoted with the thinnest bar (intron 1: hg19 chr2:176982306-17698368).

**Figure 6.**
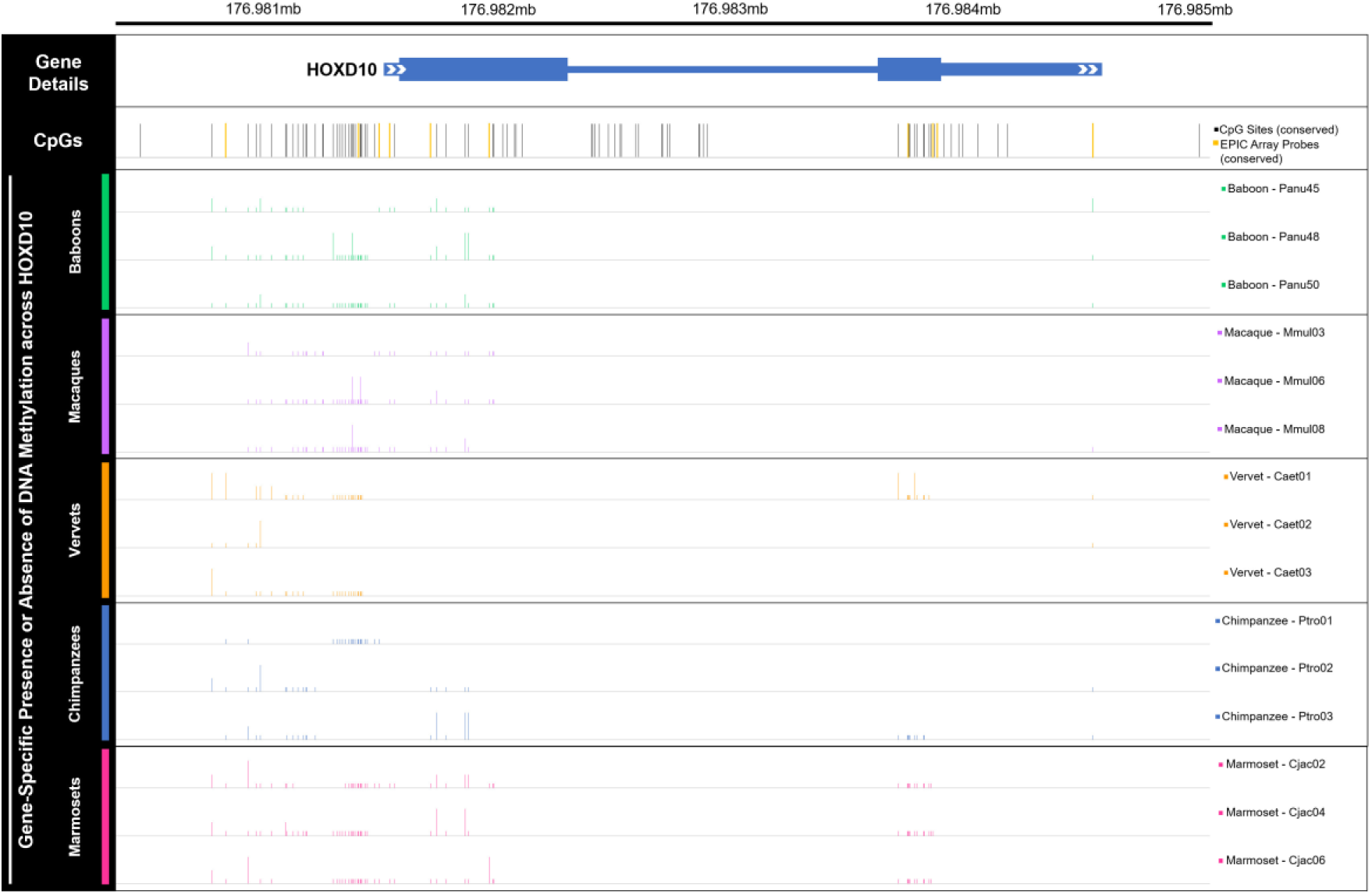
Gene-Specific Methylation Levels Across *HOXD10* in Nonhuman Primates. Bar plot of DNA methylation across the *HOXD10* gene (hg19 chr2:176981492-176984670), as well as upstream and downstream several hundred bases (hg19 chr2:176980532-176985117). Bars depict the presence (tall bar), partial presence (medium bar), or absence (low bar) of methylation at human derived CpG sites in 15 nonhuman primate samples – 3 baboons, 3 macaques, 3 vervets, 3 chimpanzees, and 3 marmosets. While regular sequencing was very successful, bisulfite sequencing was less successful, with several sequence reads uninterpretable. As such, nonhuman primate methylation data is only available for a subset of the CpGs known in humans. Partial presence of methylation was called when sequencing fluorescence peaks for cytosine and thymine were both present at a particular site and one was at least half the size of the other. Overall, these data provide additional information regarding gene-specific methylation levels across *HOXD10*. CpG sites that were also targeted by the EPIC array are highlighted in yellow and include cg18115040 (chr2, position 176981328), cg25371634 (chr2, position 176981422), cg13217260 (chr2, position 176981469), cg03918304 (chr2, position 176981654), cg17489939 (chr2, position 176981919), cg26708100 (chr2, position 176983815), cg10393811 (chr2, position 176983927), cg08992581 (chr2, position 176983949), and cg06005169 (chr2, position 176984634). See Table S18 and Files S8-S9 for additional information.

## Discussion

This study explored skeletal gene regulation and its relation to complex traits and species differences by evaluating genome-wide and gene-specific DNA methylation patterns in trabecular bone from several NHP species. These species are phylogenetically representative of the order and display anatomical and locomotor differences. Specifically, while the sampled cercopithecoids (baboons, macaques, vervets) display varying levels of terrestrial and arboreal quadrupedal locomotion, the sampled ape species (chimpanzees) is a knuckle-walking quadruped that also vertical climbs and occasionally walks bipedally (Cawthon Lang, 2006). Even more distinct are the sampled platyrrhines (marmosets) which locomote via vertical clinging and leaping (Cawthon Lang, 2005). Lemurs and lorises are the only major taxonomic primate groups not examined in this study. Regardless, the current sample set provided a unique opportunity to examine skeletal epigenetic differences in relation to bone morphology at an evolutionary scale. In this research we assume that adult bone methylation patterns reflect gene regulation that enables the maintenance of post-maturation skeletal morphologies. Some of these patterns may be impacted by environmental forces, like diet and biomechanical loading, but we attempted to control for these variables by only including NHPs from captive colonies with similar environmental exposures. Nevertheless, we are unable to ascertain how substantially the adult skeletal methylation in our sample set deviates from the regulatory patterns that initially arose during early skeletal element development. Despite these limitations, the novel data collected in this study can help to inform the relationship between epigenetic variation and skeletal traits. First, we evaluated the association between intra-specific variation in skeletal DNA methylation and intra-specific variation in femur morphology. Second, we assessed inter-specific DNA methylation differences and the potential contribution of these lineage-specific patterns to species-specific morphologies.

### Intra-specific DMPs are not readily associated with morphological variants

With respect to intra-specific morphology, very few sites were found to be differentially methylated. DMPs were only identified in association with baboon bicondylar femur length, baboon maximum femur length, macaque proximal femur width, macaque medial condyle width, vervet superior shaft width, vervet inferior shaft width, vervet anatomical neck height, and chimpanzee anatomical neck length (Table S8). Additionally, most of these associations only identified one DMP. This limited number of associations may be due to the small amount of variation identified in each morphology. This point is supported in that almost all the morphologies with methylation associations also have the highest intra-specific variation in size (Figure S5, Files S2-S3). Further, because the animals included in this study were born and raised in captivity, their limited exposure to environmental variation may limit both the range of phenotypic variation and methylation variation present within species. Alternatively, the limited number of associations may simply be due to the small sample sizes within each species. Increasing the number of individuals included in this study would improve the power to detect morphology-related DMPs. Indeed, for our largest NHP sample set (baboons: n=28, 189,858 CpG sites evaluated), there is only 11% power to detect a 10% change in methylation (Δβ = 0.1) between two groups given an FDR cutoff of 0.05 (Graw, Henn, Thompson, & Koestler, 2019). Further, larger sample sizes might also reveal larger amounts of variation within each morphology. Thus, investigations employing larger sample sets will provide more insight on these fronts. Nevertheless, it may be the case that DNA methylation variation does not have a large influence on nonpathological femur morphology within NHPs.

Of the few intra-specific methylation patterns associated with morphology, they likely have weak functional effects. Research has shown that individual site-specific methylation changes are not readily associated with differential gene expression (Bork et al., 2010; Chen et al., 2011; Koch et al., 2011). Rather, differential gene expression is made possible through the accumulation of several methylation changes within promotor regions (Suzuki & Bird, 2008) or across the gene body (Singer, Kosti, Pachter, & Mandel-Gutfreund, 2015). Detecting accumulations of methylation changes across genes should have been possible in the current study as 4-19 probes targeted each gene on average (Table S4). Thus, the individual sites identified in this study are not due to experimental limitations and likely have limited impacts on gene regulation. This is further supported by the degree of methylation variation observed among these DMPs. Sites that have an average change in mean methylation less than 10% (Δβ < 0.1) are thought to have little biological relevance (Hernando-Herraez et al., 2013). Thus, those DMPs with small changes in methylation likely have little to no biological function. Further, some of the DMPs with Δβ ≥ 0.1 appear to be highly influenced by small subsets of the sample sets. For example, in the case of macaque proximal width, the majority of methylation change is due to two individuals (1 female and 1 male) that have extremely low methylation at cg19349877 as compared to other macaques (File S3). Additionally, in the case of chimpanzee anatomical neck length, the 273 DMPs identified are largely affected by one individual that has a longer anatomical neck than other chimpanzees. Lastly, the limited enrichment of GO biological processes and KEGG pathway functions among detected DMPs indicates a lack of common function among these associated regions. Overall, these finding suggest that while some DMPs appear to be associated with intra-specific morphological variation in NHPs, not enough evidence is present to support them having a functional role in the development and maintenance of this morphological variation.

### Differential methylation is observed among NHP species

With respect to inter-specific variation, several sites were found to have significant species-specific methylation differences. Specifically, out of the 39,802 sites examined, 650 species-specific DMPs were identified in baboons, 257 in macaques, 639 in vervets, 2,796 in chimpanzees, and 13,778 in marmosets, and these span 7,320 genes (Tables S11-S12). These numbers of species-specific DMPs did not substantially change when accounting for age scaled to life expectancy instead of exact age (File S4). However, many of these DMPs had biologically insignificant changes in mean methylation. Thus, several Δβ cutoff thresholds were considered – from a 10% change in mean methylation (Δβ ≥ 0.1) up to a 40% change in mean methylation (Δβ ≥ 0.4) – which reduced the overall numbers of species-specific DMPs. Regardless of Δβ cutoff threshold, more species-specific DMPs were found in the platyrrhine marmosets, followed by the great ape chimpanzees, and lastly the cercopithecoid baboons, macaques, and vervets. This trend in numbers of species-specific DMPs is expected given the known phylogenetic relationships between primates, with cercopithecoids more closely related to apes and platyrrhines more distantly related to both groups (Perelman et al., 2011; Rogers & Gibbs, 2014). However, the number of marmoset-specific DMPs is substantially larger than those for other taxa. While this discrepancy may reflect marmosets having more species-specific changes, aspects of the experimental design may also contribute to it. Marmosets are the only platyrrhine included in this study, so although some marmoset-specific DMPs are truly specific to marmosets, others may be specific to all platyrrhines. Comparably, cercopithecoid-specific DMPs and catarrhine-specific DMPs may be cancelled out from this study, as such changes would be shared between all cercopithecoids and catarrhines, respectively. Second, since marmosets are the most phylogenetically distant from humans as compared to the other NHPs included in this study (Perelman et al., 2011; Rogers & Gibbs, 2014), the probe filtering steps may have biased downstream data in favor of finding significant results primarily in marmosets. Third, the marmoset data itself has a slightly different normalized distribution, with more mean methylation levels of 50%, than that for other NHPs (Figure S4), which may be due to the chimerism often observed in marmosets. Levels of chimerism in marmoset skeletal tissues have not been examined. However, blood-derived tissues are known to have high levels of chimerism (>50% chimeric), as compared to epithelial tissues which have the lowest levels of all tissues examined thus far (12% chimeric) (Malukiewicz et al., 2015; Ross, French, & Orti, 2007). Even if trabecular bone has low chimerism, its close proximity to blood cells in bone marrow increases the chance of highly chimeric cell contamination. Additionally, although when assessed computationally, the filtered array probes examined in this study appear to hybridize sufficiently for marmosets (Figure S3), there may be other unknown biological or technical issues that impede proper DNA methylation analyses from the EPIC array in marmosets and may have inflated the overall number of marmoset-specific DMPs identified. Given the disproportionate number of marmoset-specific DMPs identified and the unusual nature of these data, interpreting the biological relevance of these data requires caution.

The number of species-specific DMPs identified in this study, is comparable to those identified in a previous study that assessed methylation patterns in blood from chimpanzees, bonobos, gorillas, and orangutans using the 450K array and similar alignment criteria filtering methods with a focus on sites that had a Δβ ≥ 0.1 (Hernando-Herraez et al., 2013). This research used a final set of 99,919 probes that were shared across all great ape species and covered 12,593 genes with at least 2 probes per gene. Out of these, 2,284 species-specific DMPs were found in humans, 1,245 in chimpanzees and bonobos, 1,374 in gorillas, and 5,501 in orangutans (Hernando-Herraez et al., 2013). In the current study, when a Δβ ≥ 0.1 threshold was used, only 1,572 chimpanzee-specific DMPs were identified (Table S11). This number is lower than the number found in previous research comparing chimpanzees to other great apes (Hernando-Herraez et al., 2013), species that are evolutionarily closer to chimpanzees than are cercopithecoids and platyrrhines (Perelman et al., 2011; Rogers & Gibbs, 2014). However, the total number of sites examined in the present study is approximately one-third of that examined in the prior great ape study. Thus, a 3-fold increase in the number of sites examined might identify a 3-fold increase in chimpanzee-specific DMPs as compared to cercopithecoids and platyrrhines, which would be closer to expectations (Hernando-Herraez et al., 2013).

Additionally, the number of DMPs that distinguish species from one another in the current study is substantially smaller than the numbers of DMPs identified between different skeletal tissue types, between different age cohorts, and between individuals with different skeletal disease states within a NHP species (Housman et al., 2018). This finding is to be expected since differences in DNA methylation which regulate gene expression (Singer et al., 2015; Suzuki & Bird, 2008) should increase or decrease with the degree of differences between cellular functions among comparative groups (Zhang et al., 2013). In the case of different tissues, substantial DNA methylation differences likely promote distinct gene regulation that is necessary for cells in different tissues to promote different functions. In the case of different age cohorts, slightly fewer DNA methylation differences within the same skeletal tissue may allow cells to emphasize efforts on growth and development in juveniles, as compared to maintenance in adults, without altering the general skeletal-related functions of this tissue. In the case of osteoarthritic disease states, even fewer regulatory changes may be needed to initiate the dysregulation of tissue function. Finally, the comparatively small number of DNA methylation differences between adult, skeletally healthy, NHP species seems reasonable given that other studies have also noted the presence of more regulatory variation within species than between species (Uebbing et al., 2016). In summary, epigenetic and regulatory differences, which control and enable age-dependent organ functions, should be greater within a species when comparing between tissue types, age cohorts, and disease states, than when comparing between species.

When evaluating how well the methylation patterns at species-specific DMPs cluster samples, a Δβ ≥ 0.4 threshold is necessary to achieve clustering into distinct species (Figure 3). Less stringent Δβ thresholds are able to separate apes, platyrrhines, and cercopithecoids into distinct groups, but the cercopithecoids do not form monophyletic species groups at these thresholds (Figure S6). Global changes in methylation show similar phylogenetic patterns. While average species methylation patterns reveal a well-supported tree topology that reflects known phylogenetic relationships between taxa (Perelman et al., 2011; Rogers & Gibbs, 2014) (Figure 4), global changes in methylation among individual animals do not distinguish cercopithecoids into distinct monophyletic groups (Figure S7). As before, parsing down these global changes to only species-specific DMPs with Δβ ≥ 0.4 fixes the phylogeny (Figure S8). In previous studies, global methylation changes were able to separate great apes into species-specific phylogenetic groups (Hernando-Herraez et al., 2013). Conversely, the need for a higher Δβ cutoff threshold to distinguish species in the current study may be due to evolutionary reasons.

The divergence times between cercopithecoids (baboons, macaques, and vervets) are comparable to those between great apes (humans, chimpanzees, gorillas, and orangutans) (Perelman et al., 2011). Thus, divergence times do not explain why global changes in methylation are unable to resolve species-specific phylogenetic clades in the current study as compared to previous research. On the other hand, while nonhuman great apes have experienced higher rates of molecular evolution as compared to humans (Elango, Thomas, NISC Comparative Sequencing Program, & Yi, 2006), baboons and macaques have slow rates of molecular evolution as compared to other cercopithecoids (Elango, Lee, Peng, Loh, & Yi, 2009), which may correspond to slower rates of epigenetic evolution. This might make baboons and macaques appear more similar to vervets than expected, and further, this may make resolving the phylogenetic divergences between cercopithecoids more difficult than that between the great apes. Additionally, the number of sites included in the present study (39,802) as compared to previous studies (99,919) (Hernando-Herraez et al., 2013) may limit the ability of the present study to fully resolve species-specific lineages. This is reinforced by the low support of several branches in the phylogeny based on global changes in methylation across individual animals (Figure S7). Conversely, sample size was likely not a contributing factor to the discrepancies between the current study (n=58) and previous studies (n=32) (Hernando-Herraez et al., 2013), as the current study has a slightly larger sample size. However, the number of individuals per species was more uniform in previous studies (Hernando-Herraez et al., 2013) than in the current study.

Alternatively, the fact that global changes in methylation are unable to fully resolve cercopithecoid species-specific phylogenetic clades in the current study, may instead indicate that not enough time has passed for cercopithecoid species to evolve fixed epigenetic changes between taxa in this tissue. Additionally, it is possible that epigenetic variation at many of the sites examined in this study are under balancing selection in cercopithecoids which prevents these markers from accurately resolving the evolutionary divergences between these species. Previous research of gene regulation differences between species has found that some deviations in gene expression may be under directional or balancing selection (Romero, Ruvinsky, & Gilad, 2012; Whitehead & Crawford, 2006). Given the relatively low variance in methylation across species in combination with lineage-specific changes in mean methylation at the species-specific DMPs identified in this study, it is possible that these loci are experiencing similar selective pressures (Romero et al., 2012). Nevertheless, most inter-specific regulatory differences appear to be under stabilizing selection or neutral evolution (Brawand et al., 2011; Gilad, 2012; Romero et al., 2012), and this cannot be ruled out in the present study.

### Inter-specific DMPs are found in genes enriched for functions associated with skeletal traits

The evolution of methylation changes along specific NHP lineages is associated with several functions that may contribute to species-specific phenotypic differences (Files S6-S7). First, several skeletal tissue functions are enriched in species-specific DMPs. Among almost all NHPs, cellular adhesion functions are highly enriched. Cellular adhesion is necessary for cells to attach to other cells or extracellular matrix, which is a necessity for bone cells (Mbalaviele, Shin, & Civitelli, 2006). Additionally, in baboons, vervets, chimpanzees, and marmosets, anatomical developmental processes are enriched, and in chimpanzees and marmosets, pattern specification processes, limb development, and skeletal system development are enriched. Lastly, in marmosets, specific skeletal functions, such as osteoblast differentiation and ossification are also enriched. Overall, these functions validate that most patterns of differential methylation relate to skeletal tissue function, regulation, development, and maintenance, as well as to larger anatomical developmental processes. Additionally, functions not specific to the skeletal system were identified. In chimpanzees and marmosets, transcription and gene expression regulatory functions were enriched. Further, in marmosets, functions related to the development of skeletal muscles, nerves, the brain, the heart, blood vessels, kidneys, eyes, and ears were also enriched. All together, these findings suggest that many species-specific changes in methylation may contribute to the regulation of complex phenotypic changes. While this relationship was not observed for intra-specific skeletal morphology variation, other skeletal traits not examined in this study may be related. Nevertheless, many of the genes associated with the described functions only contain an average of 1-2 differentially methylated sites. As described above, individual site-specific methylation changes are not readily associated with differential gene expression (Bork et al., 2010; Chen et al., 2011; Koch et al., 2011). Therefore, the enriched functions identified are likely not true biological effects due to methylation differences on their own. Rather they hint at biological effects that may be the result of the combined effects of several genetic, epigenetic, and other regulatory processes.

Finally, some of the genes containing species-specific DMPs overlap with those previously identified as being differentially methylated in other tissues among primates. Specifically, the *HOXD* cluster which is involved in limb development shows differential methylation among species. In the current study, *HOXD9* shows species-specific hypermethylation at 1 site in baboons and species-specific hypomethylation at 2 sites in chimpanzees, while in previous work, *HOXD9* shows hypermethylation in archaic hominins as compared to modern humans (Gokhman et al., 2014). In the *HOXD* cluster, *HOXD10* also contains interesting inter-specific DNA methylation patterns (Figure 5, Table S13), has an active role in anatomical development, and has been found to be differentially methylated among hominins (Gokhman et al. 2014). Thus, it was selected for subsequent DNA methylation profiling and analysis at a higher resolution using gene-specific sequencing techniques.

*HOXD10* specifically codes for a protein that functions as a sequence-specific transcription factor which is expressed in the developing limb buds and is involved in differentiation and limb development. In the current study, each NHP shows low to intermediate methylation levels across the gene body (Figure 5, Table S13). A similar pattern is observed in the gene-specific methylation data, which further reveals that *HOXD10* is not highly methylated in NHPs. However, across all taxa, some clusters of hypermethylation are found upstream of the gene and at the start of the gene body, with marmosets on average displaying more methylation in the gene body than other taxa (Figure 6, Table S18). The consistency between bisulfite sequence and EPIC array data in marmosets does not clarify the unusual nature of the marmoset EPIC array data (Figure S4), so more data are needed in these samples to better characterize marmoset *HOXD10* methylation patterns. In the *HOXD10* gene body of hominins, humans display hypomethylation, Neandertals displayed intermediate methylation levels, and Denisovans displayed high levels of methylation (Gokhman et al., 2014). The variation of methylation patterns in this gene body suggest that intermediate methylation levels may be a more ancestral epigenetic state for this region in the primate lineage, while the extreme hypermethylation of this region in Denisovans and the extreme hypomethylation of this region in humans may be derived epigenetic states. Previous work has proposed that methylation differences in *HOXD10* may be associated with phenotypic distinctions between modern human and archaic hominin limbs (Gokhman et al., 2014). While the current study did not find substantial associations between methylation variation and aspects of femur morphology within NHP species, further work to understand the role of differential methylation of *HOXD10* in promoting morphological changes of the limb should be explored.

### Conclusions and future directions

In conclusion, while only a few significant associations were identified between methylation and femur morphologies, several significant differences in methylation were observed inter-specifically. Moreover, these species-specific DMPs were found in genes enriched for functions associated with complex skeletal traits. This is the first study to characterize DNA methylation patterns in skeletal tissues from a taxonomically diverse set of NHPs, and it is the first study to directly compare these patterns to the nonpathological morphologies of the skeletal elements from which the tissues were derived. This design enabled an initial exploration of skeletal gene regulation and its relation to complex traits and species differences for which little else is currently known.

Ultimately, this work forms a foundation for future explorations of gene regulation and skeletal trait evolution in primates. The dataset collected and investigated here provides an initial sampling of skeletal methylation patterns in NHPs, and future primate skeletal epigenetics investigations can add to these data at population- and evolutionary-scales. At a population-level, increasing the number of individuals from one species can provide further insight into how genetic variation interacts with gene regulation to contribute to a wider variety of healthy and pathological skeletal traits. Additionally, sampling individuals of different ages would provide further insight into how gene regulation across development contributes to skeletal anatomy. Adult bone methylation patterns are limited in their ability to fully inform how gene regulation contributes to the differential development of skeletal morphologies. In the present study, we assume that regulatory patterns present in adult skeletal tissues are involved in (or at least not inhibiting) the maintenance of adult skeletal morphology, and that the regulatory patterns that enable the maintenance of adult skeletal traits reflect those that enabled the development of those traits. Future work in the field of primate skeletal epigenetics could explicitly test the degree to which adult bone methylation patterns reflect patterns that arise during early skeletal element development. Lastly, at an evolutionary-level, increasing the number of represented species, both extant and extinct, will further inform our understanding of gene regulation evolution in primates and how patterns of variation contribute to skeletal phenotypes.

## Acknowledgements

This work was supported by the National Institutes of Health (P01HL028972 to Anthony G. Comuzzie, P40OD010965 to Matthew J. Jorgensen, P40RR019963 to Jay Kaplan); the Leakey Foundation (Research Grant for Doctoral Students to G.H.); the Wenner-Gren Foundation (Gr. 9310 to G.H.); the Nacey Maggioncalda Foundation (James F. Nacey Fellowship to G.H.); the International Primatological Society (to G.H.); Sigma Xi (Grant-in-Aid of Research to G.H.); the ASU Center for Evolution and Medicine (Venture Fund to G.H.); the ASU Graduate Research and Support Program (to G.H.). Additionally, this investigation used resources that were supported by the Southwest National Primate Research Center grant P51OD011133 from the Office of Research Infrastructure Programs, National Institutes of Health.

We thank Eric D. Johnson and members of the Department of Genetics at the Texas Biomedical Research Institute, including Anthony G. Comuzzie, Anne Sheldrake, Jaydee Foster, Kara Peterson, Mel Carless, and Laura Cox, for helpful discussions. We also thank Megann Phillips for assistance with PCR primer design.

Newly reported data have been made available on NCBI’s Gene Expression Omnibus and are accessible through the GEO SuperSeries accession number GSE103332, which includes the following SubSeries accession numbers: GSE103279 (intra-specific baboon DNA methylation data), GSE103271 (intra-specific macaque DNA methylation data), GSE103280 (intra-specific vervet DNA methylation data), GSE94677 (intra-specific chimp DNA methylation data), GSE103328 (intra-specific marmoset DNA methylation data), and GSE103287 (inter-specific DNA methylation data).

