## Supplemental Text for "Intra- and Inter-Specific Investigations of Skeletal DNA Methylation and Femur Morphology in Primates"

**Supplemental Materials**

**Supplemental Text**

**Additional Details on Genome-Wide DNA Methylation Profiling Methods**

Trabecular bone is used in this study because several human skeletal epigenetic studies are based on trabecular bone, and it is important to standardize tissue type for comparative purposes. Trabecular bone comprises the internal spongy osseous tissue that contributes to femoral shape morphology and remodeling, which begins before birth and continues throughout life (Clarke, 2008). Trabecular bone in growing individuals influences both trabecular and cortical morphology in adulthood (Wang, Ghasem-Zadeh, Wang, Iuliano-Burns, & Seeman, 2011), and this suggests that the epigenetics of trabecular bone may be of more interest than that of cortical bone. Lastly, although trabecular bone is not ideal for epigenetic analyses because it contains several cell types (Horvath et al., 2015), statistical methods can correct for this heterogeneity.

Tissues were collected from the same portion of the femur in order to minimize potential variation between samples and comparative groups. Cartilage methylation patterns are known to vary between joints and between different sites within a joint (den Hollander et al., 2014; Jeffries et al., 2016; Loughlin & Reynard, 2015; Moazedi-Fuerst et al., 2014; Rushton et al., 2014). Although similar studies of bone methylation patterns have not been conducted yet, the number and types of cells, and therefore epigenetic signatures, are expected to vary across different portions of the femur. Additionally, regulatory patterns in skeletal tissues may be impacted by environmental forces, like diet and biomechanical loading, as well as growth and developmental processes. We attempted to control for environmental variables by only including NHPs from captive colonies with similar environmental exposures and by collecting tissues from the same portion of the femur, which further ensured consistency in the effects of biomechanical loading across individuals and species. We controlled for the potential effects of growth and developmental processes by sampling adult NHPs that have reached sexual maturity and are maintaining skeletal morphology, as opposed to developing and growing it.

Illumina Infinium MethylationEPIC microarrays (EPIC arrays) analyze the methylation status of over 850,000 sites throughout the genome, covering over 90% of the sites on the Infinium HumanMethylation450 BeadChip, as well as an additional 350,000 sites within enhancer regions. For each sample, 400ng of genomic DNA was bisulfite converted using the EZ DNA Methylation$^{TM}$ Gold Kit according to the manufacturer's instructions (Zymo Research), with modifications described in the Infinium Methylation Assay Protocol. Following manufacturer guidelines (Illumina), this processed DNA was then whole-genome amplified, enzymatically fragmented, hybridized to the arrays, and imaged using the Illumina iScan system.

**Additional Details on Methylation Data Processing**

Raw fluorescent data were normalized to account for the noise inherent within and between the arrays themselves. Specifically, we performed a normal-exponential out-of-band (Noob) background correction method with dye-bias normalization (Triche, Weisenberger, Van Den Berg, Laird, & Siegmund, 2013) to adjust for background fluorescence and dye-based biases. We followed this with a between-array normalization method (functional normalization) (Fortin et al., 2014) which removes unwanted variation by regressing out variability explained by the control probes present on the array as implemented in the minfi package in R (Aryee et al., 2014; Fortin, Triche, & Hansen, 2016) which is part of the Bioconductor project (Huber et al., 2015). This method has been found to outperform other existing approaches for studies that compare conditions with known large-scale differences (Fortin et al., 2014), such as those assessed in this study.

After normalization, methylation values (β values) for each site were calculated as the ratio of methylated probe signal intensity to the sum of both methylated and unmethylated probe signal intensities (Equation 1). These β values range from 0 to 1 and represent the average methylation levels at each site across the entire population of cells from which DNA was extracted (0 =completely unmethylated sites, 1 = fully methylated sites). Every β value in the Infinium platform is accompanied by a detection p-value, and those with failed detection levels (p-value > 0.05) in greater than 10% of samples were removed from subsequent analyses. Additionally, samples in which more than 30% of the β value had a detection p-value > 0.05 were removed from downstream analyses. Because β values have high heteroscedasticity, they are not statistically valid for use in differential methylation analyses (Du et al., 2010). Thus, M values were calculated as the log transformed ratio of methylated signal to unmethylated signal and used in the statistical analyses described below.

The probes on the arrays were designed to hybridize specifically with human DNA, so our use of nonhuman primate (NHP) DNA required that probes non-specific to any of the included NHP genomes, which could produce biased methylation measurements, be computationally filtered out and excluded from downstream analyses. This was accomplished using two different methods modified from (Hernando-Herraez et al., 2013; Ong et al., 2014). For both methods, we used blastn (Altschul et al., 1997) to map the 866,837 50bp probes onto the baboon, macaque, vervet, chimpanzee, and marmoset genomes (Table S2) using an e-value threshold of $e^{-10}$. Probes that successfully mapped to each genome, had only 1 unique BLAST hit, and targeted CpG sites were retained. This resulted in 39% of all probes being retained for baboons, 39% for macaques, 39% for vervets, 76% for chimpanzees, and 17% for marmosets (Table S3). These proportions were slightly lower than expected based on the previous findings of 44% retention for baboons using the 450K array (Housman, Havill, Quillen, Comuzzie, & Stone, 2018) and of 61% retention for Cynomologus macaques using the 450K array (Ong et al., 2014). The altered design of the EPIC array as compared to the 450K array may explain both discrepancies. Additionally, the higher quality of the Cynomologus macaque genome (Assembly: Macaca_fascicularis_5.0; Accession: GCF_000264685.2; Average Scaffold Length: 88,649,475; Average Contig Length: 86,040) as compared to the baboon, Rhesus macaque, and vervet genomes used in this study (Table S2), may explain the latter discrepancy. Regardless, and as expected, species more closely related to humans (e.g., chimpanzees) have higher numbers of reliably mapped EPIC array probes than species more distantly related to humans (e.g., marmosets).

Subsequent *in silico* analyses based on sequence alignment criteria (Hernando-Herraez et al., 2013) and based on gene symbol criteria (Ong et al., 2014) were then used on the mapped EPIC array probes. For the first filtering method (alignment filter criteria), probes were only retained if they had 0 mismatches in the 5bp closest to and including the CpG site and if they had 0-2 mismatches in the 45bp not including the CpG site. This resulted in 24% of all probe being retained for baboons, 24% for macaques, 24% for vervets, 71% for chimpanzees, and 9% for marmosets (Figure S1, Table S3). Conversely, for the second filtering method (gene symbol filter criteria), we identified the closest NHP gene to each probe site and checked for corresponding gene name matches between humans and each NHP. This information was obtained from different sources for each taxon (Table S2). Only those probes with partial or complete gene matches were retained. This resulted in 22% of all probes being retained for baboons, 19% for macaques, 23% for vervets, 32% for chimpanzees, and 10% for marmosets (Figure S1, Table S3). Overall, 39,802 probes are shared among all taxa for the alignment filter criteria, while 36,248 probes are shared among all taxa for the gene symbol filter criteria.

Overall, similar number of probes were retained for both filtering criteria, with the exception of macaque and chimpanzee probe sets. This discrepancy is due to the lack of gene information from Ensembl BioMart for these genome versions (Table S2). Again, species more closely related to humans have higher numbers of retained probes than species more distantly related to humans. Additionally, in chimpanzees, the proportion of probes retained using the alignment criteria (72%) exactly matches the proportion previously found using the same filtering methods for the 450K array (Hernando-Herraez et al., 2013). Furthermore, these retained probes maintained wide and comparable distributions throughout the genome. Depending on the species, the retained probes cover between 9,779 and 24,107 genes, with between 4 and 19 probes per gene, and they are spread across several genomic locations and proximities to CpG islands (Table S4). Nevertheless, between the probes retained using the alignment filter criteria and the probes retained using the gene symbol criteria, there is only partial overlap. Specifically, about half of the resulting probes for each filtering technique overlapped with one another for baboons, macaques, vervets, and marmosets, and for chimpanzees, almost all of the retained probes using the gene symbol criteria overlapped with those retained using the alignment filter criteria (Figure S2). This discrepancy is likely due to the incomplete nature of each NHP genome annotation, as described above. Given the general lack of overlap between filtered probe sets, it was necessary to determine which filtering method was more optimal for subsequent analyses.

We accomplished this by determining which filtered probe set more effectively measured DNA methylation in the DNA of each species. First, we used the detection p-value of each probe as an assessment of that probe's hybridization efficiency. We performed Spearman correlation tests and found that the hybridization efficiency of each probe is significantly correlated with the alignment quality of each probe to each NHP genome, and thus, the degree of sequence conservation (Table S5). The majority of filtered probes for both *in silico* methods passed quality controls and produced robust signals on the array, indicating that either filtering technique may be appropriate for future research. However, because probes retained using the alignment filter criteria had a higher proportion of successfully hybridized probes than the probes retained using the gene symbol filter criteria (Figure S3) and because the alignment filter criteria are less influenced by the degree of genome assembly annotation, using the alignment filter criteria likely produces more reliable results. Thus, only these data were used for downstream differential methylation analyses.

Lastly, cross-reactive probes (Chen et al., 2013), probes containing SNPs at the CpG site, probes detecting SNP information, probes detecting methylation at non-CpG sites, and probes targeting sites within the sex chromosomes were filtered out. Using the alignment filter criteria, this resulted in a final set of 189,858 probes for baboons, 190,898 probes for macaques, 191,639 probes for vervets, 576,804 probes for chimpanzees, 68,709 probes for marmosets, and 39,802 probes shared among species (Figure S4). Conversely, using the gene symbol filter criteria, this resulted in a final set of 165,529 probes for baboons, 146,585 probes for macaques, 175,592 probes for vervets, 254,231 probes for chimpanzees, 75,002 probes for marmosets, and 33,254 probes shared among species (Figure S4).

Lastly, to ensure that probes showing significant methylation differences were not biased by known NHP SNPs, we also assessed NHP SNPs overlaps. Specifically, for probes that successfully mapped to a NHP genome, we determined how many NHP SNPs (Table S2) overlapped with the entire probe region and whether any overlapping SNPs were located at the targeted CpG site (File S1). Because the quality of SNP data is variable across species and because only a small number of CpG-SNP overlaps were identified in each alignment filter probe set (1,158 probes for baboons, 82 probes for macaques, 121 probes for vervets, 299 probes for chimpanzees, 731 probes for marmosets, and 406 probes shared among species), we did not filter out probes containing CpG-SNP overlaps before running statistical analyses. However, for the small number of CpG-SNP overlaps that were identified in significantly differentially methylated probes, we note these probes as such in corresponding supplementary files (Table S9, FileS3-S5).

**Additional Details on Statistical Analysis of Differential Methylation**

We designed and tested general linear models (GLMs) which related variables of interest to the DNA methylation patterns for each site, while accounting for other fixed effects such as sex, age, steady state weight, batch effects, and latent variables. The specific GLM design matrices used in this study are outlined below.

Latent variables were calculated using the iteratively re-weighted least squares approach in the sva function of the sva package in R (Jaffe & Irizarry, 2014; Leek, Johnson, Parker, Jaffe, & Storey, 2012; Leek & Storey, 2007, 2008), and these were used in the GLMs to account for cell heterogeneity. Alternative methods to account for cell heterogeneity exist, but they are specific to whole blood (Jaffe & Irizarry, 2014; Morris & Beck, 2015), require reference epigenetic data, or are reference free methods (Houseman, Molitor, & Marsit, 2014) that are comparable to the sva method (Kaushal, Zhang, Karmaus, & Wang, 2015). Out of the known cell types in skeletal tissues (Horvath et al., 2015), only chondrocytes and osteoblasts have reference epigenomes available on the International Human Epigenomics Consortium, and these are only for humans, not NHPs. Thus, because no standard method is available to correct for the heterogeneous cell structure in NHP skeletal tissue, we chose the described sva method.

Each GLM design matrix was fit to corresponding M value array data by generalized least squares using the lmFit function of the limma package in R (Huber et al., 2015; Phipson, Lee, Majewski, Alexander, & Smyth, 2016; Ritchie et al., 2015), and the estimated coefficients and standard errors for each morphology were computed. For each coefficient, an empirical Bayes approach was applied using the eBayes function in the limma package in R (Lönnstedt & Speed, 2002; McCarthy & Smyth, 2009; Phipson et al., 2016; Smyth, 2004) to compute moderated t-statistics, log-odds ratios of differential methylation, and associated p-values adjusted for multiple testing (Benjamini & Hochberg, 1995).

GLM Design Matrices

Intra-specific analyses in baboons:

methylation ~ femur morphology + sex + age + weight + latent variables

Intra-specific analyses in macaques, vervets, and marmosets when $n > 5$:

methylation ~ femur morphology + sex + age + latent variables

Intra-specific analyses in chimpanzees and marmosets when $n \leq 5$:

methylation ~ femur morphology + latent variables

Inter-specific analyses:

methylation ~ taxonomic group + sex + age + batch effects + latent variables

### Supplemental Figures

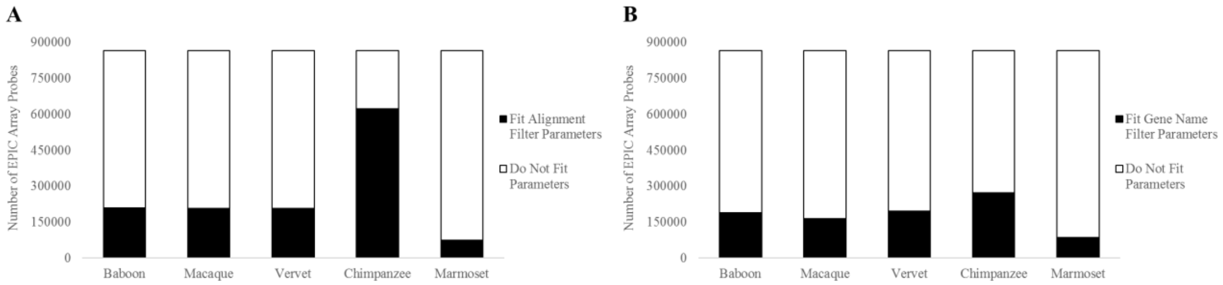

**Figure S1. Filtering Effects on EPIC Array Probes for NHPs.**
Bar chart showing the percent of EPIC array probes retained for each species using the alignment filter criteria (A) or using the gene symbol filter criteria (B). Using the alignment filter criteria, 24% of probes are retained for baboons, 24% for macaques, 24% for vervets, 72% for chimpanzees, and 9% for marmosets. Using the gene filter criteria, 22% of probes are retained for baboons, 19% for macaques, 23% for vervets, 32% for chimpanzees, and 10% for marmosets. See Tables S2-S4, and File S1 for additional details.

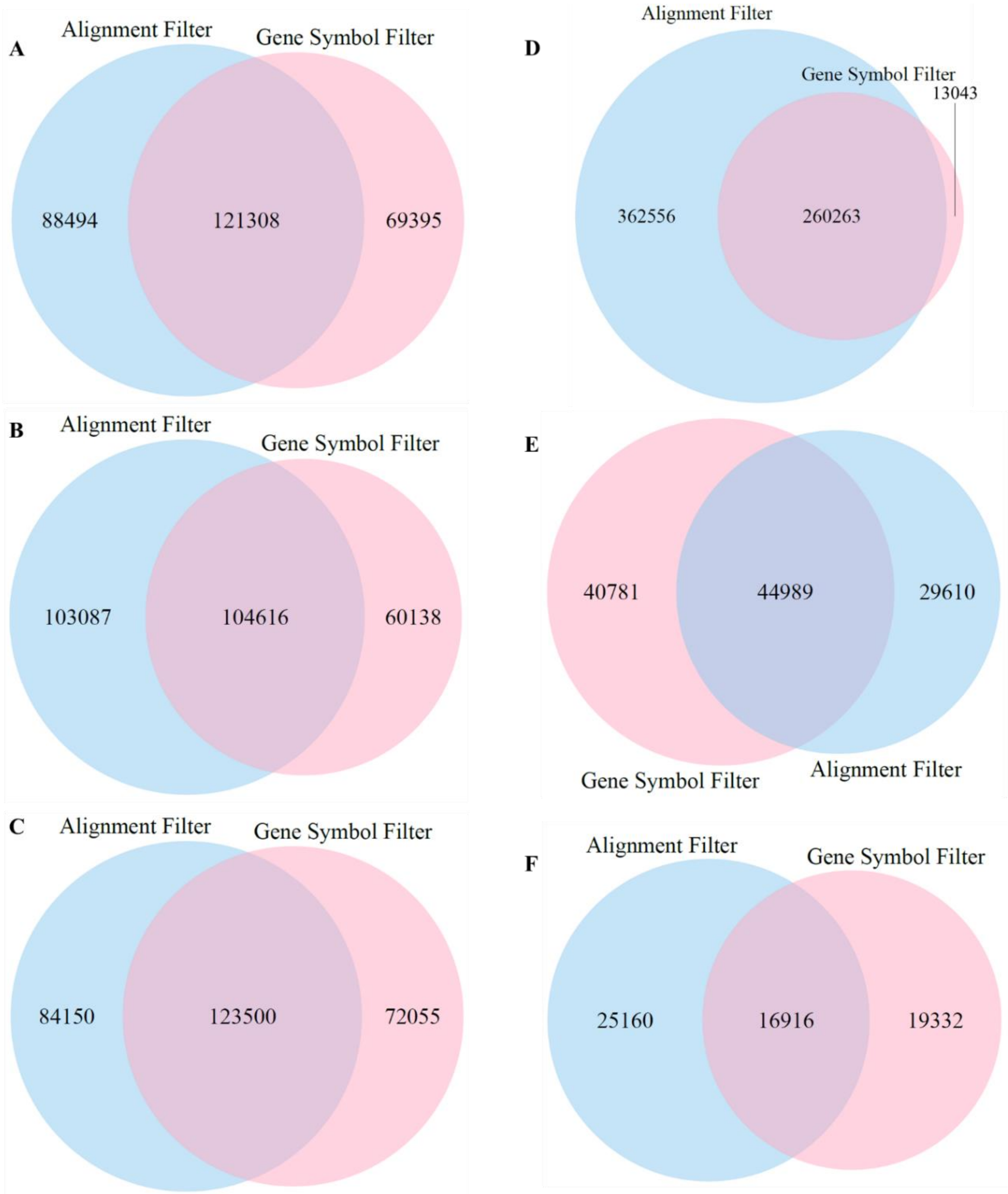

**Figure S2. Overlap of EPIC Array Probes for NHPs Using Different Filtering Methods.**
Venn diagrams showing the number of probes that overlap between the alignment filter criteria and the gene symbol criteria for each species. (A) For baboons, out of the 209,802 probes that meet the alignment filter criteria and the 190,703 probes that meet the gene symbol criteria, 121,308 probes (58% and 64% respectively) overlap in both filters. (B) For macaques, out of the 207,703 probes that meet the alignment filter criteria and the 164,754 probes that meet the gene symbol criteria, 104,616 probes (50% and 63% respectively) overlap in both filters. (C) For vervets, out of the 207,650 probes that meet the alignment filter criteria and the 195,555 probes that meet the gene symbol criteria, 123,500 probes (59% and 63% respectively) overlap in both filters. (D) For chimpanzees, out of the 622,819 probes that meet the alignment filter criteria and the 273,306 probes that meet the gene symbol criteria, 260,263 probes (42% and 95% respectively) overlap in both filters. (E) For marmosets, out of the 74,599 probes that meet the alignment filter criteria and the 85,770 probes that meet the gene symbol criteria, 44,989 probes (60% and 52% respectively) overlap in both filters. (F) For probes that align to all NHP genomes, out of the 42,076 probes that meet the alignment filter criteria and the 36,248 probes that meet the gene symbol criteria, 16,916 probes (40% and 47% respectively) overlap in both filters.

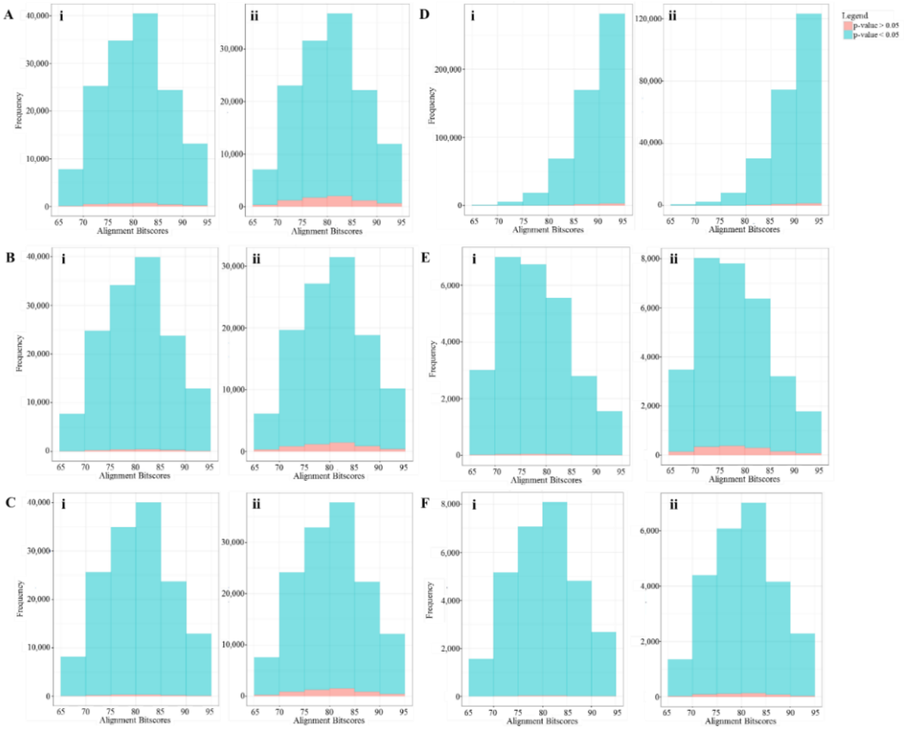

**Figure S3. Hybridization Efficiencies of EPIC Array Probes Retained for NHPs.**
Histogram of alignment bitscores for EPIC array probes with detection p-values > 0.05 (red) and < 0.05 (blue). These p-values were averaged across all samples within each species, and probes included meet (i) the alignment filter criteria or (ii) the gene symbol filter criteria. (A) Baboons: For probes meeting the alignment filter criteria (i), 3,880 had detection p-values > 0.05, and 205,922 had detection p-values < 0.05. For probes meeting the gene symbol filter criteria (ii), 10,571 had detection p-values > 0.05, and 180,132 had detection p-values < 0.05. For all probes that successfully mapped to the baboon genome with e-values < $e^{-10}$, 337,818 had only unique BLAST hits and targeted a CpG site, 21,977 had detection p-values > 0.05, and 315,841 had detection p-values < 0.05. (B) Macaques: For probes meeting the alignment filter criteria (i), 2,586 had detection p-values > 0.05, and 205,117 had detection p-values < 0.05. For probes meeting the gene symbol filter criteria (ii), 7,442 had detection p-values > 0.05, and 157,312 had detection p-values < 0.05. For all probes that successfully mapped to the macaque genome with e-values < $e^{-10}$, 335,046 had only unique BLAST hits and targeted a CpG site, 17,821 had detection p-values > 0.05, and 317,225 had detection p-values < 0.05. (C) Vervets: For probes meeting the alignment filter criteria (i) 2,007 had detection p-values > 0.05, and 205,643 had detection p-values < 0.05. For probes meeting the gene symbol filter criteria (ii), 7,732 had detection p-values > 0.05, and 187,823 had detection p-values < 0.05. For all probes that successfully mapped to the vervet genome with e-values < $e^{-10}$, 336,786 had only unique BLAST hits and targeted a CpG site, 15,405 had detection p-values > 0.05, and 321,381 had detection p-values < 0.05. (D) Chimpanzees: For probes meeting the alignment filter criteria (i), 6,120 had detection p-values > 0.05, and 616,699 had detection p-values < 0.05. For probes meeting the gene symbol filter criteria (ii), 3,241 had detection p-values > 0.05, and 270,065 had detection p-values < 0.05. For all probes that successfully mapped to the chimpanzee genome with e-values < $e^{-10}$, 657,913 had only unique BLAST hits and targeted a CpG site, 9,982 had detection p-values > 0.05, and 647,931 had detection p-values < 0.05. (E) Marmosets: For probes meeting the alignment filter criteria (i), 595 had detection p-values > 0.05, and 74,004 had detection p-values < 0.05. For probes meeting the gene symbol filter criteria (ii), 3,993 had detection p-values > 0.05, and 81,777 had detection p-values < 0.05. For all probes that successfully mapped to the marmoset genome with e-values < $e^{-10}$, 143,407 had only unique BLAST hits, and targeted a CpG site, 7,481 had detection p-values > 0.05, and 135,926 had detection p-values < 0.05. (F) All NHP Species Combined: For probes meeting the alignment filter criteria (i), 201 had detection p-values > 0.05, and 41,875 had detection p-values < 0.05. For probes meeting the gene symbol filter criteria (ii), 770 had detection p-values > 0.05, and 35,478 had detection p-values < 0.05.

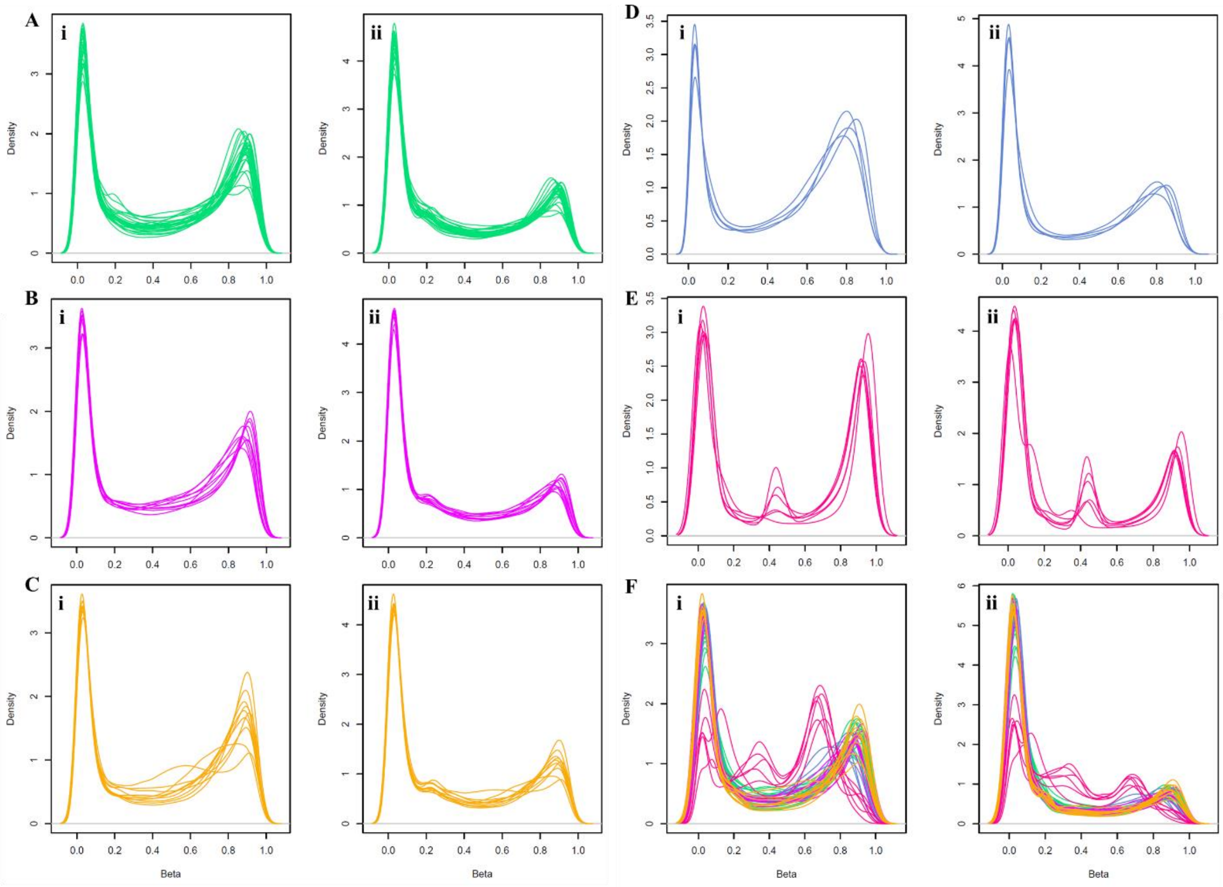

**Figure S4. Normalized and Filtered Methylation Data.**
Density plots of β-values after normalization and probe filtering using the alignment criteria (i) or gene symbol criteria (ii) for baboons (A), macaques (B), vervets (C), chimpanzees (D), marmosets (E), and all the combination of all taxa together (F), respectively.

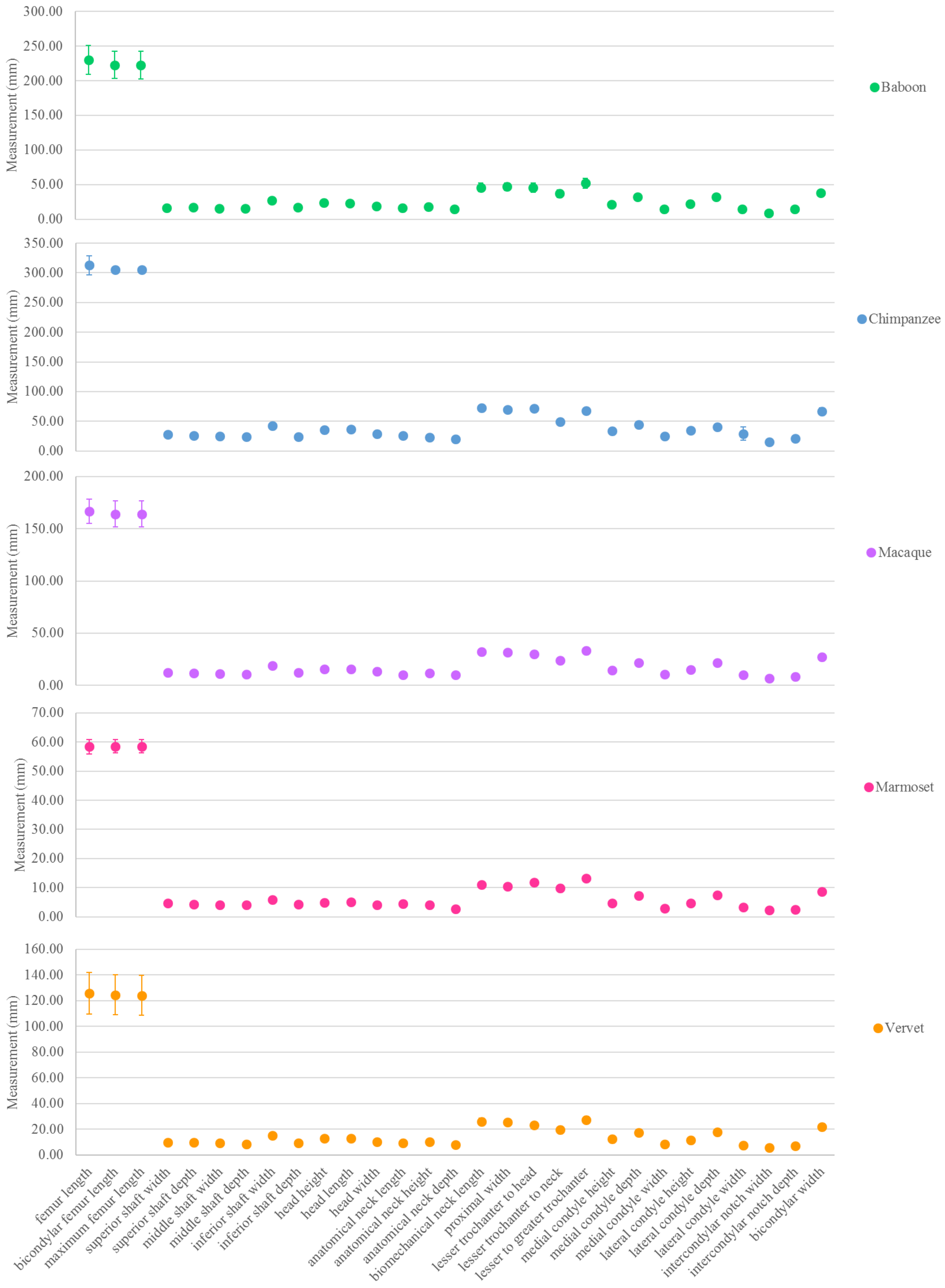

**Figure S5. Results of NHP Morphological Measurements.**
Plot of linear morphological measurements in each species. Plot depicts the average measurement in millimeters with error bars displaying one standard deviation in each direction. See File S2 for additional details.

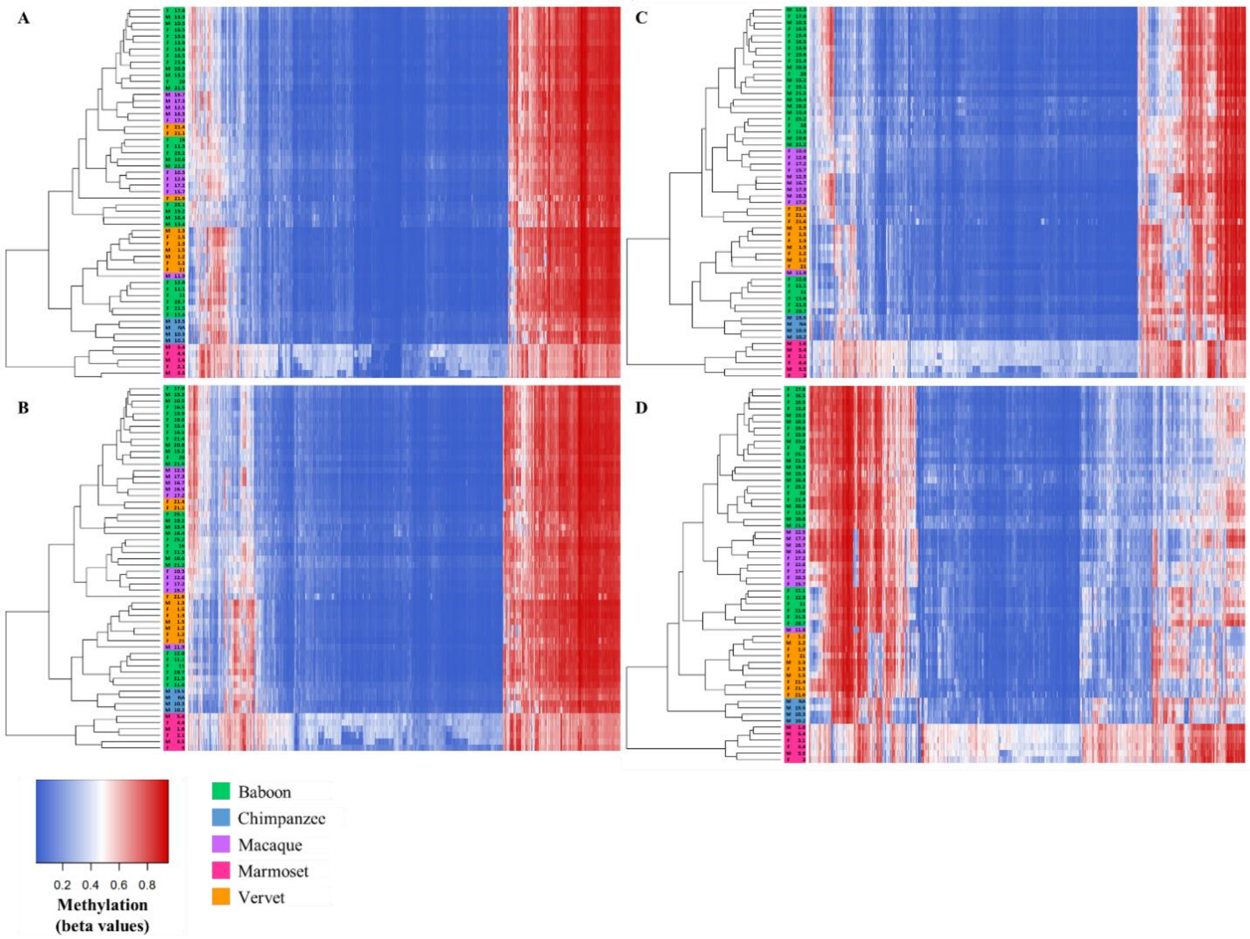

**Figure S6. Methylation Levels at Species-Specific DMPs with Various Δβ Threshold Cutoffs Identified in the Inter-Specific Study.**

Heatmap depicting (A) the DNA methylation levels (β values) of all species-specific DMPs (x-axis) in all NHP samples (n=58), (B) the DNA methylation levels (β values) of all species-specific DMPs with average absolute Δβ values greater than 0.1 between each taxonomic group (x-axis) in all NHP samples (n=58), (C) the DNA methylation levels (β values) of all species-specific DMPs with average absolute Δβ values greater than 0.2 between each taxonomic group (x-axis) in all NHP samples (n=58), and (D) the DNA methylation levels (β values) of all species-specific DMPs with average absolute Δβ values greater than 0.3 between each taxonomic group (x-axis) in all NHP samples (n=58). The sex and age of each NHP are also provided (y-axis). Red indicates higher methylation at a DMP, while blue indicates lower methylation at a DMP. The dendrogram of all samples (y-axis) clusters individuals based on the similarity of their methylation patterns. Samples cluster into the large taxonomic groupings of platyrrhines, cercopithecoids, and apes, but cercopithecoids do not cluster by species for any of these filtering levels. See File S4 for additional details.

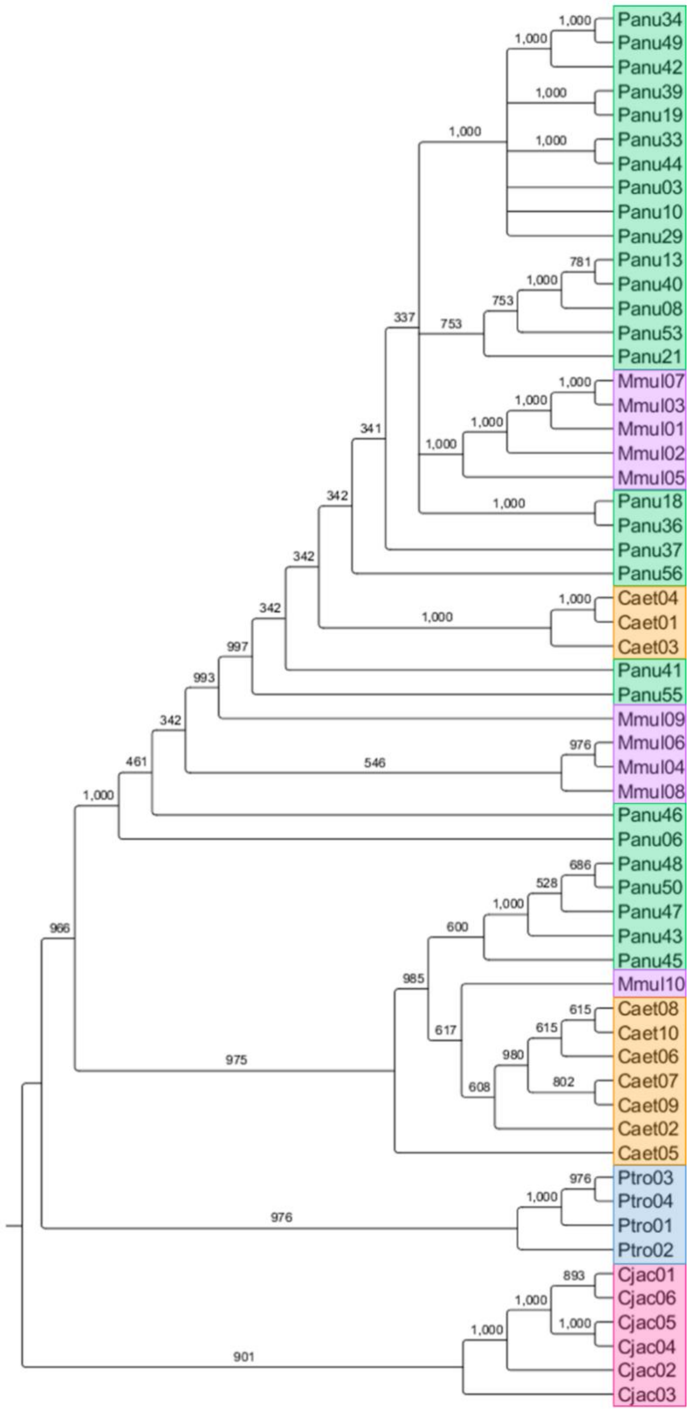

**Figure S7. Phylogeny Based on Individual-Level Global Changes in Methylation.**
Observed phylogenetic relationship among NHPs when considering individual-level global changes in methylation. This tree was constructed using the methylation levels for all of the finalized 39,802 filtered probes. We used Euclidean distances to calculate the difference between every two individuals, and estimated a neighbor joining tree using this distance matrix. For the resulting tree, 1000 bootstraps were performed to determine confidence values for each branch. The number provide at each node indicates the number of bootstrap replicates that support it out of the 1000 performed.

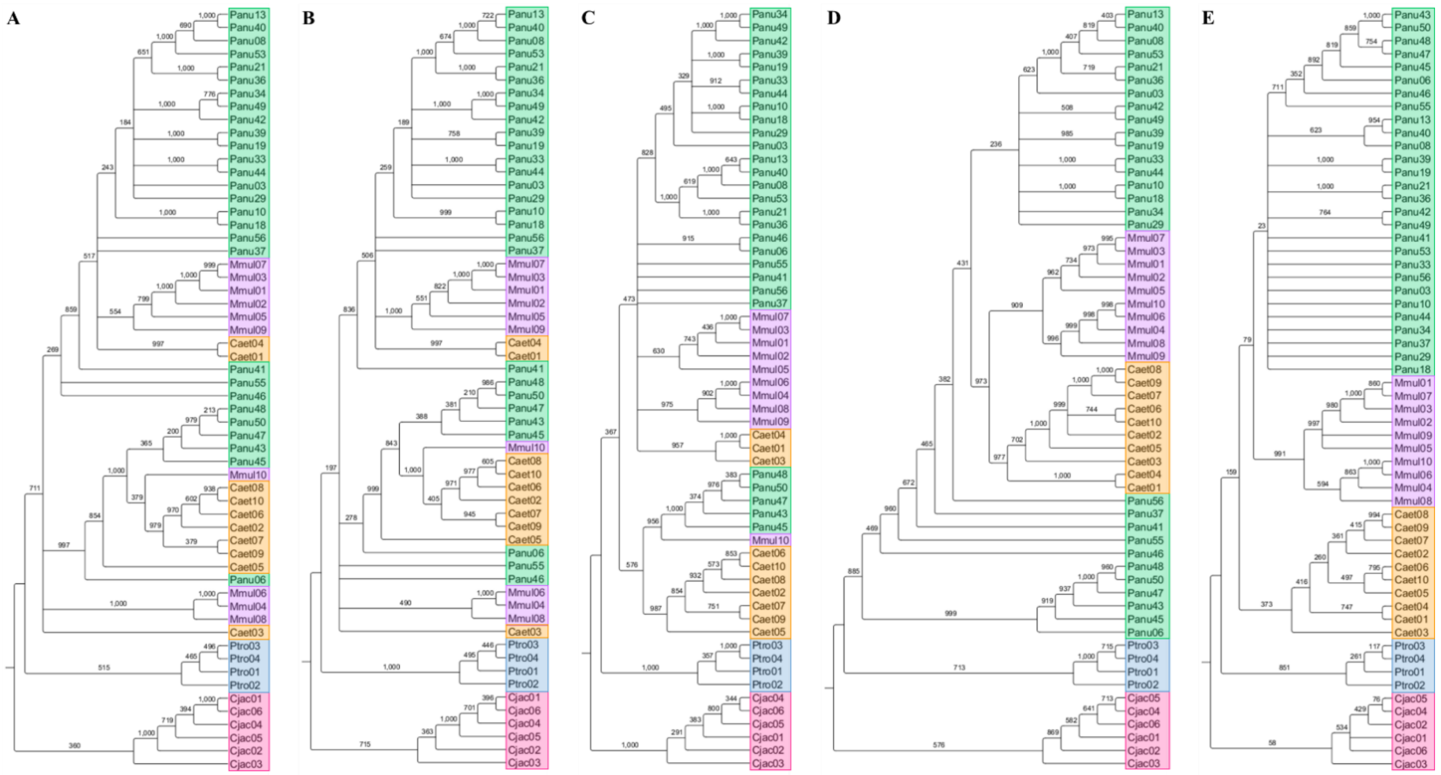

**Figure S8. Phylogeny Based on Individual-Level Changes in Methylation at Species-Specific DMPs with Various Δβ Threshold Cutoffs.**

Observed phylogenetic relationship among NHPs when considering individual-level global changes in methylation. Trees were constructed using the methylation levels for (A) all species-specific DMPs, (B) species-specific DMPs with average absolute Δβ values greater than 0.1 between each taxonomic group, (C) species-specific DMPs with average absolute Δβ values greater than 0.2 between each taxonomic group, (D) species-specific DMPs with average absolute Δβ values greater than 0.3 between each taxonomic group, and (E) species-specific DMPs with average absolute Δβ values greater than 0.4 between each taxonomic group. We used Euclidean distances to calculate the difference between every two individuals, and estimated a neighbor joining tree using this distance matrix. For the resulting tree, 1000 bootstraps were performed to determine confidence values for each branch. The number provide at each node indicates the number of bootstrap replicates that support it out of the 1000 performed.

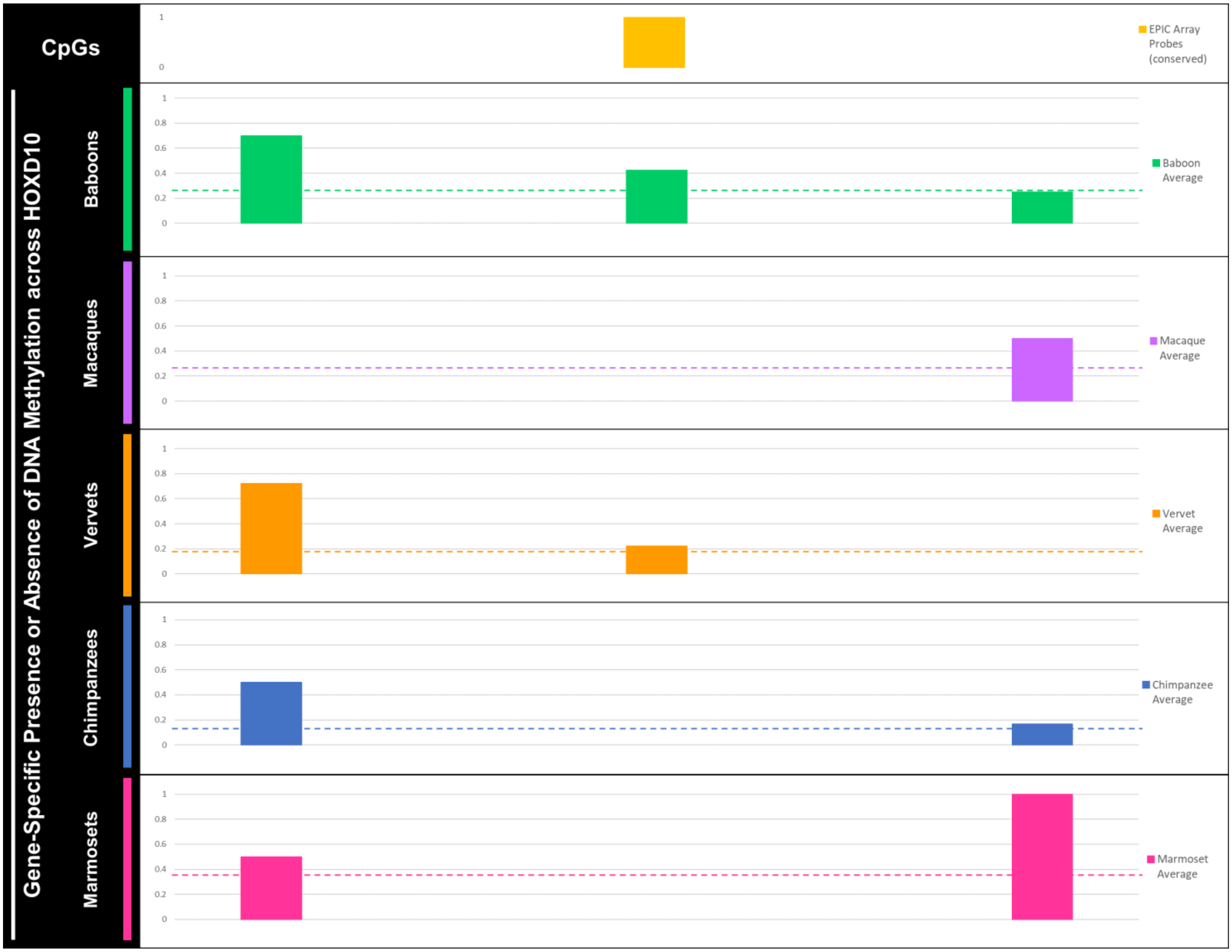

**Figure S9. Gene-Specific Methylation Levels at a Test Locus.**

Bar plot of DNA methylation across three CpG sites in the *HOXD10* gene (hg19 chr2:176981492-176984670) that are conserved among all NHPs sampled. The middle CpG site corresponds to cg02193236 (chr2, position 176980734) which was also targeted on the EPIC array. The left-most CpG site is the first conserved CpG site just upstream of cg02193236, and the right-most CpG site is the first conserved CpG site just downstream of cg02193236. Bars display the average methylation level for each NHP species (baboon, macaque, vervet, chimpanzee, and marmoset) as calculated using the gene-specific methylation data (presence = 1, absence = 0, partial methylation = 0.5) collected from all samples. While regular sequencing resulted in successful calling of nucleotide bases in sequence reads, bisulfite sequencing resulted in sequence reads with higher numbers of bases unable to be called. Specifically, no methylation data were available for the first two CpG sites in macaques. Dotted lines represent the average methylation levels of the cg02193236 test locus for each NHP species as determined using the EPIC array. The cg02193236 probe had appropriate detection levels in all species and is a conserved CpG site in all sampled species but was filtered out during data processing and so was not included in further genome-wide associate analyses. The methylation levels of cg02193236 as determined using the EPIC array and the methylation levels around this test locus as determined using gene-specific sequencing are similar. These findings supplement previous methylation array replication studies that have found data to be highly correlated in replicate chimpanzee tissue samples (>98% correlated) (Pai, Bell, Marioni, Pritchard, & Gilad, 2011) and in replicate macaque tissue samples (97% correlated) (Ong et al., 2014). Combining these lines of evidence, we believe our EPIC array data are reliable. Nevertheless, additional data collections may aid in confirming this further. See Table S1, Table S19, and Files S10-S11.

**Supplemental Table Legends**
All supplemental tables are provided in TableS1-S19.xlsx.

**Table S1. NHP Sample Set.**
Details of NHP sample set, outlining each sample's animal identification number (Animal ID), species (Species), sex (Sex), age in years (Age), life expectancy estimate in years (Life Expectancy) (Rowe, 1996), age scaled to estimated life expectancy (Scaled Age), the identification number of the beadchip (Array ID) and array (Position) that each sample was run on, and whether downstream gene-specific analyses of the test locus or the loci across the *HOXD10* gene were performed on a sample (Gene-Specific Analyses).

**Table S2. NHP Genomes Used for Probe Filtering Methods.**
Baboon, macaque, vervet, chimpanzee, and marmoset genome assemblies and accession numbers used for probe filtering methods. Average scaffold lengths and average contig lengths for each genome also provided, as well as the gene information data sources used in the gene symbol probe filtering method and the SNP data sources.

**Table S3. Number of EPIC Array Probes Retained for NHPs.**
The numbers of probes that successfully mapped to each NHP genome with e-values less than e-10, had only unique BLAST hits, and targeted a CpG site (Total Mapped Probes), probes that fit the alignment filter criteria (Alignment Filter Probes), and probes that fit the gene symbol filter criteria (Gene Symbol Filter Probes).

**Table S4. Genomic Distribution of EPIC Array Probes Retained for NHPs.**
Details on the gene associations, genomic locations, and proximities to CpG islands (based on the human genome hg19) of all probes that successfully mapped to each NHP genome with e-values less than e-10, had only unique BLAST hits, and targeted a CpG site (Total Mapped Probes), probes that fit the alignment filter criteria (Alignment Filter Probes), and probes that fit the gene symbol filter criteria (Gene Symbol Filter Probes). Number of Genes indicates the number of unique gene symbols associated with probes. Probes Per Gene indicates the average number of probes that target each associated gene. For genomic locations, TSS 200 and TSS 1500 indicate the transcription start site areas between the start of the gene and 200bp upstream or 1500bp upstream respectively, 5'UTR and 3'UTR indicate the untranslated regions of genes, 1st Exon indicates the first exon within a gene, ExonBnd indicates the gene body exons, and Gene Body indicates any area within the exons and introns of a gene. For proximity to CpG islands, Island indicates areas within CpG islands, North Shelf indicates areas 2-4kb upstream from a CpG island, North Shore indicates areas up to 2kb upstream from a CpG island, South Shelf indicates areas 2-4kb downstream from a CpG island, South Shore indicates areas up to 2kb downstream from a CpG island, and Open Sea indicates isolated CpG site in the genome.

**Table S5. Alignment Parameter Correlations of EPIC Array Probes Retained for NHPs.**
Results of Spearman correlation tests between EPIC array probe detection p-values and alignment quality parameters. These parameters included the alignment bitscores, e-values, and percent identity. For all probes that successfully mapped to one of the NHP genomes with e-values < e-10, had only unique BLAST hits, and targeted a CpG site (Total Mapped Probes), probes that fit the alignment filter criteria (Alignment Filter Probes), and probes that fit the gene symbol filter criteria (Gene Symbol Filter Probes), significant correlations were identified for alignment bitscores, e-values, and percent identities.

**Table S6. NHP Morphological Measurements.**
Linear morphological measurements collected from the right femur of each NHP. Measurements selected were based on (McHenry & Corruccini, 1978; Terzidis et al., 2012). See Figure 2 for a visual representation of these measurements.

**Table S7. Samples and Latent Variables Included in Intra-Specific Study.**
The number of samples and latent variables included in each GLMM testing for associations between DNA methylation levels and femur morphology within each species.

**Table S8. Number of Significant DMPs Identified in the Intra-Specific Study.**
Table showing the number of significant DMPs associated with each linear morphology in each species. Results are shown for probes filtered using the alignment criteria. The number of total sites tested for baboons was 189,858, for macaques was 190,898, for vervets was 191,639, for chimpanzees was 576,804, and for marmosets was 68,709. The number of samples and latent variables included in each GLMM are provided in Table S7.

**Table S9. Gene Details of Significant DMPs Identified in the Intra-Specific Study.**
Table showing details on all significant DMPs associated with a linear morphology in a NHP species. Table includes the identification number of each significant DMP probe (EPIC Array Probe ID), as well as the log fold difference in M values between each comparative group (Log Fold Change in M Values), the p-values for each DMP after accounting for multiple testing (Adjusted P-Value), and additional annotation information for each significant DMP probe (Human Gene Symbol, Baboon Gene Symbol, Baboon Chromosome, Baboon CpG Position).
*One probe (cg07644869) contains a potential SNP variant at the CpG site in chimpanzees.

**Table S10. KEGG Pathways Enriched for Significant DMPs Associated with Anatomical Neck Lengths in Chimpanzees.**
KEGG pathways that are significantly enriched (FDR $< 0.05$) for significant DMPs that are associated with anatomical neck lengths in chimpanzees, taking into account the differing number of probes per gene present on the EPIC array. Table includes the identification numbers (KEGG ID) and pathways (KEGG Pathway) for each significantly enriched KEGG term, the total number of genes associated with each KEGG pathway (No. Genes Total), the number of genes with significant DMPs that are also associated with each KEGG pathway (No. Gene with DMPs), the p-value for over-representation of each KEGG pathway (P-Value), and the false discovery rate for each KEGG pathway (FDR).

**Table S11. Number of Significant Species-Specific DMPs Identified.**
Details on the number of species-specific DMPs identified for each NHP species in total, when the average absolute change in $\beta$ values between each pairwise species comparison ($\Delta\beta$) is greater than 0.1, greater than 0.2, greater than 0.3, or greater than 0.4. The pattern of methylation (hypermethylated, hypomethylated, or a mixture of both) of each species-specific DMP is also indicated.

**Table S12. Genomic Distribution of Significant Species-Specific DMPs Identified.**
Details on the gene associations, genomic locations, and proximity to CpG islands (based on the human genome hg19) of all probes that successfully mapped to each NHP genome with e-values less than e-10, had only unique BLAST hits, targeted a CpG site, and fit the alignment filter criteria. Number of Genes indicates the number of unique gene symbols associated with DMPs. Probes Per Gene indicates the average number of DMPs within each associated gene. For genomic locations, TSS 200 and TSS 1500 indicate the transcription start site areas between the start of the gene and 200bp upstream or 1500bp upstream respectively, 5'UTR and 3'UTR indicate the untranslated regions of genes, 1st Exon indicates the first exon within a gene, ExonBnd indicates the gene body exons, and Gene Body indicates any area within the exons and introns of a gene. For proximity to CpG islands, Island indicates areas within CpG islands, North Shelf indicates areas 2-4kb upstream from a CpG island, North Shore indicates areas up to 2kb upstream from a CpG island, South Shelf indicates areas 2-4kb downstream from a CpG island, South Shore indicates areas up to 2kb downstream from a CpG island, and Open Sea indicates isolated CpG site in the genome.

**Table S13. DMP Analysis Results for *HOXD10* Gene.**
Table describing the differentially methylated position (DMP) analysis results for the *HOXD10* genes. Table includes the identification number of each DMP examined (EPIC Array Probe ID), as well as additional annotation information for each of these sites (Gene Symbols, Chromosomes, CpG Positions), whether each site was found to be significantly differentially methylated between taxonomic groups and if so whether the specified species was hypermethylated, hypomethylated, or had a mixture of hyper- and hypo-methylation as compared to other species (Species-Specific DMP), and the average β values for each taxonomic group (Average β Values). Of the 5 species-specific DMPs in the *HOXD10* gene of marmosets, 4 have Δβ between 0.2 and 0.3 (bold) and 1 has a Δβ < 0.1 (not bolded). Human information came from hg19.

**Table S14. NHP Gene-Specific *HOXD10* Primers.**
Primers used for the gene-specific amplification of the *HOXD10* gene using regular and bisulfite treated DNA. Details include the name of the *HOXD10* region being amplified (PCR Name), whether the primer was designed to amplify regular or bisulfite treated DNA (Primer Type), the species in which the primers were designed to amplify each *HOXD10* region (Species), the sequences for the forward and reverse primers in the 5' to 3' direction (Forward Primer Sequence (5'-3'), Reverse Primer Sequence (5'-3'), the length of amplicons in base pairs (Amplicon Length), and the optimized annealing temperature (Optimized Tm).

**Table S15. PCR Assay Specifications.**
PCR amplification temperature conditions used for both regular and bisulfite treated DNA. The annealing temperature varied for primer pairs. See optimized temperature for each primer pair in Table S14.

**Table S16. PCR Reagent Details for Regular DNA.**
Reagent initial concentrations ([initial]), final concentrations ([final]), and amounts (Volume (uL)) used for PCR amplification of regular DNA. The minimum initial concentration for DNA was 5ng/uL.

**Table S17. PCR Reagent Details for Bisulfite Treated DNA.**
Reagent initial concentrations ([initial]), final concentrations ([final]), and amounts (Volume (uL)) used for PCR amplification of bisulfite treated DNA. The minimum initial concentration for bisulfite treated DNA was 33ng/uL.

**Table S18. Gene-Specific Sequencing Results for Loci Across the *HOXD10* Gene.**
Table describing the results for the gene-specific regular and bisulfite sequencing of loci across the *HOXD10* gene (hg19 chr2:176981492-176984670), as well as upstream and downstream several hundred bases (hg19 chr2:176980532-176985117). Table includes the positions of human derived CpG sites in the *HOXD10* gene (Human CpG Position). These positions are based on the sequence alignments produced in this study (File S7-S8). CpG sites that are conserved across all sampled species are noted (Conserved in NHPs), and those targeted by the EPIC array have probe information provided (EPIC Array Probe ID). For each NHP sample examined, the presence or absence of nucleotide mutations at each CpG site is indicated (Mutation). The dinucleotides resulting from mutations are shown in parentheses, as well. These data were determined based on regular sequences. Additionally, the presence or absence of methylation (Methylation) at each CpG sites was determined based on bisulfite sequences. Abbreviations: no data collected (-), data quality too poor to call base pair (NA), methylation present at site indicated by cytosine in bisulfite sequence (yes), no methylation at site indicated by a lack of cytosine at site (no), partial methylation indicated by a partial cytosine signal at site (partial), CpG site 1bp upstream or downstream of that in humans (*), and CpG site 2bp upstream or downstream of that in humans (**). See Files S8-S9 for additional details.

**Table S19. Gene-Specific Sequencing Results for Test Locus in the *HOXD10* Gene.**
Table describing the results for the gene-specific regular and bisulfite sequencing of a test locus in the *HOXD10* gene (hg19 chr2:176981492-176984670). Table includes the positions of CpG sites in the *HOXD10* gene that are conserved across all sampled species (Human CpG Position). These positions are based on the sequence alignments produced in this study (File S10-S11). CpG sites that were also targeted by the EPIC array have probe information provided (EPIC Array Probe ID). For each NHP sample examined, the presence or absence of nucleotide mutations at each CpG site is indicated (Mutation). These data were determined based on regular sequences. Additionally, the presence or absence of methylation (Methylation) at each CpG site was determined based on bisulfite sequences. Abbreviations: no data collected (-), data quality too poor to call base pair (NA), methylation present at site indicated by cytosine in bisulfite sequence (yes), no methylation at site indicated by a lack of cytosine at site (no), partial methylation indicated by a partial cytosine signal at site (partial), CpG site 1bp upstream or downstream of that in humans (*), and CpG site 2bp upstream or downstream of that in humans (**). See Files S10-S11 for additional details.

**Supplemental File Descriptions**
All supplemental files are provided in FileS1-S11.zip. This can be found in the *bioRxiv* preprint of this paper.

**FileS1_ProbeAnnotationFilters.zip**
Nonhuman primate EPIC array probe annotation files which contain detailed information on whether each probe on the EPIC array maps to the baboon, chimpanzee, macaque, marmoset, or vervet genome, respectively, and if so, how well each probe and the mapped region align, whether associated gene symbols for humans and each nonhuman primate match, and whether nonhuman primate SNPs overlap with probes and CpGs. A probe annotation file which contains descriptions for the column headers in each probe annotation file is also provided.

**FileS2_Morphology.xlsx**
Details on the nonhuman primate morphological measurements. Linear morphological measurements used in GLMMs (Measurements), as well as triplicate measurements of linear morphologies and error calculations from these (Error). All measurements have units of millimeters. NA indicates that measurement could not be accurately collected. Error was calculated in each taxonomic group and equals the mean absolute difference divided by the mean (Corner, Lele, & Richtsmeier, 1992; White & Folkens, 2000). Only measurements with an error <5% were included in downstream analyses. This included all measurements except for intercondylar notch depth for macaques. NA indicates that measurement could not be accurately collected.

**FileS3_MorphologyDMPs.xlsx**
Gene details of the significant DMPs identified in the intra-specific morphology study. Tables show details on all significant DMPs associated with a linear morphology in a nonhuman primate species – baboon bicondylar femur length, baboon maximum femur length, macaque proximal width, macaque medial condyle width, vervet superior shaft width, vervet inferior shaft width, vervet anatomical neck height, and chimpanzee anatomical neck length. Tables include the identification number of each significant DMP probe (EPIC Array Probe ID), as well as additional information about each sample associated with the significant DMP. Specifically, the animal identification number (Animal), sex (Sex), age in years (Age), morphological measurement in millimeters (Measurements), β value for each DMP (β Values), maximum absolute change in β values for each DMP (Max Δβ), and the realized change in β values between the nonhuman primate with the largest morphology measurement and the nonhuman primate with the smallest morphology measurement for each DMP (Realized Δβ) are provided.
*One probe (cg07644869) contains a potential SNP variant at the CpG site in chimpanzees.

**FileS4_SpeciesSpecificDMPs.xlsx**
Gene details of significant species-specific DMPs identified in the inter-specific study. Tables show details on the significant DMPs identified between taxonomic groups – baboons, macaques, vervets, chimpanzees, and marmosets – and showing species-specific methylation patterns. Tables include the identification number of each significant DMP probe (EPIC Array Probe ID), whether the significant DMP probe was identified with the GLM containing age (Model Identifying DMP: Age) or with the GLM containing age scaled to life expectancy (Model Identifying DMP: Scaled Age), additional annotation information for each significant DMP probe (Human Gene Symbol, Human Chromosome, Human CpG Position, Nonhuman Primate Gene Symbol, Nonhuman Primate Chromosome, Nonhuman Primate CpG Position), the species-specific methylation observed for the species of interest (Nonhuman Primate Specific Methylation Pattern), the average β values for each taxonomic group (Average β Values), the variance in β values for each taxonomic group (Average β Values), the difference in β values (Δβ) and p-values accounting for multiple testing from either the GLM containing age (Adjusted P-Value: Age) or the GLM containing age scaled to life expectancy (Adjusted P-Value: Scaled Age) for each pairwise inter-specific comparison made for each DMP, and the average absolute difference in β values between the species of interest and each other taxonomic group (Average Absolute Difference in β Values). The table orders are divided such that those probes with the difference in β values less than 0.1, between 0.1 and 0.2,

between 0.2 and 0.3, between 0.3 and 0.4, and greater than 0.4 are separated. Mixture indicates that pairwise DNA methylation comparisons between species have a combination of hypomethylation and hypermethylation. Human annotation information is for hg19.
*Species-specific probes that contain potential SNP variants at the CpG site in the same corresponding species (3 baboon-specific probes contain potential baboon SNPs, 1 chimp-specific probe contains a potential chimp SNP, 66 marmoset-specific probes contain potential marmoset SNPs).
†Species-specific probes that contain potential SNP variants at the CpG site in different species (3 baboon-specific probes contain potential marmoset SNPs, 3 macaque-specific probes contain potential baboon SNPs, 4 macaque-specific probes contain potential marmoset SNPs, 3 vervet-specific probes contain potential baboon SNPs, 3 vervet-specific probes contain potential marmoset SNPs, 7 chimp-specific probes contain potential baboon SNPs, 1 chimp-specific probe contains a potential macaque SNP, 2 chimp-specific probes contain potential vervet SNPs, 12 chimp-specific probes contain potential marmoset SNPs, 48 marmoset-specific probes contain potential baboon SNPs, 4 marmoset-specific probes contain potential macaque SNPs, 3 marmoset-specific probes contain potential vervet SNPs, 3 marmoset-specific probes contain potential chimp SNPs).

**FileS5_GroupSpecificDMPs.xlsx**
Gene details of significant taxonomic group-specific DMPs identified in the inter-specific study. Tables show details on the significant DMPs identified between taxonomic groups – cercopithecoids, apes, and platyrrhines – and showing group-specific methylation patterns. Tables include the identification number of each significant DMP probe (EPIC Array Probe ID), whether the significant DMP probe was identified with the GLM containing age (Model Identifying DMP: Age) or with the GLM containing age scaled to life expectancy (Model Identifying DMP: Scaled Age), additional annotation information for each significant DMP probe (Human Gene Symbol, Human Chromosome, Human CpG Position, Nonhuman Primate Gene Symbol, Nonhuman Primate Chromosome, Nonhuman Primate CpG Position), the species-specific methylation observed for the species of interest (Nonhuman Primate Specific Methylation Pattern), the average β values for each taxonomic group (Average β Values), the variance in β values for each taxonomic group (Average β Values), the difference in β values (Δβ) and p-values accounting for multiple testing  from either the GLM containing age (Adjusted P-Value: Age) or the GLM containing age scaled to life expectancy (Adjusted P-Value: Scaled Age) for each pairwise inter-specific comparison made for each DMP, and the average absolute difference in β values between the species of interest and each other taxonomic group (Average Absolute Difference in β Values). The table orders are divided such that those probes with the difference in β values less than 0.1, between 0.1 and 0.2, between 0.2 and 0.3, between 0.3 and 0.4, and greater than 0.4 are separated. Mixture indicates that pairwise DNA methylation comparisons between species have a combination of hypomethylation and hypermethylation. Human annotation information is for hg19. Also contains details on the GO biological processes and KEGG pathways enriched for significant group-specific DMPs.
*Group-specific probes that contain potential SNP variants at the CpG site in the same corresponding group (12 cercopithecoid-specific probes contain potential cercopithecoid SNPs, 1 ape-specific probe contains a potential ape SNP, 67 platyrrhine-specific probes contain potential platyrrhine SNPs).
†Group-specific probes that contain potential SNP variants at the CpG site in different groups (1 cercopithecoid-specific probe contains a potential ape SNP, 18 cercopithecoid-specific probes contain potential platyrrhine SNPs, 10 ape-specific probes contain potential cercopithecoid SNPs, 10 ape-specific probes contain potential platyrrhine SNPs, 1 ape-specific probe contains a potential SNP for both cercopithecoids and apes, 65 platyrrhine-specific probes contain potential cercopithecoid SNPs, 3 platyrrhine-specific probes contain potential ape SNPs).

**FileS6_SpeciesSpecificGO.xlsx**
GO biological processes enriched for significant species-specific DMPs identified in the inter-specific study. Tables contain the GO biological process terms that are significantly enriched (FDR < 0.05) for species-specific DMPs – baboons, macaques, vervets, chimpanzees, and marmosets – taking into account the differing number of probes per gene present on the EPIC array and assessed in the current study. Tables include the identification

numbers (GO ID) and terms (GO Biological Process Term) for each significantly enriched GO term, the total number of genes associated with each GO term (No. Genes Total), the number of genes with significant DMPs that are also associated with each GO term (No. Gene with DMPs), the p-value for over-representation of each GO term (P-Value), and the false discovery Rate for each GO term (FDR).

**FileS7_SpeciesSpecificKEGG.xlsx**
KEGG pathways enriched for significant species-specific DMPs identified in the inter-specific study. Tables contain the KEGG pathways that are significantly enriched (FDR < 0.05) for species-specific DMPs – baboons, macaques, vervets, chimpanzees, and marmosets – taking into account the differing number of probes per gene present on the EPIC array and assessed in the current study. Tables include the identification numbers (KEGG ID) and pathways (KEGG Pathway) for each significantly enriched KEGG term, the total number of genes associated with each KEGG pathway (No. Genes Total), the number of genes with significant DMPs that are also associated with each KEGG pathway (No. Gene with DMPs), the p-value for over-representation of each KEGG pathway (P-Value), and the false discovery rate for each KEGG pathway (FDR).

**FileS8_HOXD10_total_trim_qc50_align_manualfix_order_samples_names-examples.fas**
FASTA sequence alignment of the processed, regular and bisulfite gene-specific sequences of loci across *HOXD10* that were evaluated in the inter-specific study. These sequences are aligned to the regions surrounding and including *HOXD10* from several primates that were obtained from the EPO whole-genome multiple alignments of several primate genomes [Ensembl Compara.8_primates_EPO] (Paten, Herrero, Beal, Fitzgerald, & Birney, 2008; Paten, Herrero, Fitzgerald, et al., 2008). Some reference sequences were bisulfited converted *in silico* to aid in alignment and analysis. See File S9 for raw sequences.

**FileS9_HOXD10_Sequences-examples.zip**
Folder containing the raw gene-specific sequences of loci across *HOXD10* that were evaluated in the inter-specific study. Each regular and bisulfite converted sequence has an unprocessed chromatogram file (*.ab1) and a sequence file (*.seq).

**FileS10_HOXD10_total_trim_qc50_align_manualfix_order_sample_names-1and2.fas**
FASTA sequence alignment of the processed, regular and bisulfite gene-specific sequences of the test locus (cg02193236) and its surrounding region in *HOXD10.* These data were used to determine the reliability of the EPIC array in assessing DNA methylation levels. These sequences are aligned to the regions surrounding and including *HOXD10* from several primates that were obtained from the EPO whole-genome multiple alignments of several primate genomes [Ensembl Compara.8_primates_EPO] (Paten, Herrero, Beal, et al., 2008; Paten, Herrero, Fitzgerald, et al., 2008). Some reference sequences were bisulfited converted *in silico* to aid in alignment and analysis. See File S11 for raw sequences.

**FileS11_HOXD10_Sequences-1and2.zip**
Folder containing the raw gene-specific sequences of the test locus (cg02193236) and its surrounding region in *HOXD10* that were used to determine the reliability of the EPIC array in assessing DNA methylation levels. Each regular and bisulfite converted sequence has an unprocessed chromatogram file (*.ab1) and a sequence file (*.seq).
